# Accurate localization and coactivation profiles of the Frontal Eye Field and Inferior Frontal Junction: an ALE and MACM fMRI meta-analysis

**DOI:** 10.1101/2022.08.11.503474

**Authors:** Marco Bedini, Emanuele Olivetti, Paolo Avesani, Daniel Baldauf

**Affiliations:** Center for Mind/Brain Sciences (CIMeC), University of Trento, Trento, Italy; NeuroInformatics Laboratory (NILab), Bruno Kessler Foundation (FBK), Trento, Italy; Department of Psychology, University of California, San Diego, La Jolla, California, USA

**Author notes:** Corresponding author at: Center for Mind/Brain Sciences, University of Trento via delle Regole, 101, 38123, Trento (TN), Italy.

**Keywords:** Prefrontal Cortex, Visual Attention, Saccadic Eye Movements, Working memory, Activation Likelihood Estimation, Meta-analytic Connectivity Modeling

## Abstract

The frontal eye field (FEF) and the inferior frontal junction (IFJ) are prefrontal structures involved in mediating multiple aspects of goal-driven behavior. Despite being recognized as prominent nodes of the networks underlying spatial attention and oculomotor control, and working memory and cognitive control, respectively, the limited quantitative evidence on their precise localization has considerably impeded the detailed understanding of their structure and connectivity. In this study, we performed an activation likelihood estimation (ALE) fMRI meta-analysis by selecting studies that employed standard paradigms to accurately infer the localization of these regions in stereotaxic space. For the FEF, we found the highest spatial convergence of activations for prosaccades and antisaccades paradigms at the junction of the precentral sulcus and superior frontal sulcus. For the IFJ, we found consistent activations across oddball/attention, working memory, Stroop, and task-switching paradigms at the junction of the inferior precentral sulcus and inferior frontal sulcus. We related these clusters to previous meta-analyses, sulcal/gyral neuroanatomy, and a recent comprehensive brain parcellation, highlighting important differences compared to their results and taxonomy. Finally, we employed the ALE peak coordinates as seeds to perform a meta-analytic connectivity modeling (MACM) analysis, which revealed systematic coactivation patterns spanning the frontal, parietal and temporal cortices. We decoded the behavioral domains associated with these coactivations, suggesting that these may allow FEF and IFJ to support their specialized roles in flexible behavior. Our study provides meta-analytic groundwork for investigating the relationship between functional specialization and connectivity of two crucial control structures of the prefrontal cortex.

## Introduction

Owing to the capabilities that likely derive from the massive expansion in the cortical surface allowed by the folding patterns of the cortex (Van Essen 2007; Welker 1990; Zilles et al. 2013), which particularly involved the prefrontal and association cortices (Donahue et al. 2018; Toro et al. 2008), humans possess one of the most complex behavioral repertoires in nature (Mesulam 1998; Miller and Cohen 2001). A fundamental aspect of functional specialization in the human brain is its relationship with cortical neuroanatomy (Van Essen 2007). Microstructural features pertaining to cortical architecture (i.e., cyto- and myelo-architecture), such as cell types and layer organization, are a major determinant of the functional organization of the brain, and they provide important information about regional segregation (Brodmann 1909; Amunts et al. 2020). Over the past 30 years, magnetic resonance imaging (MRI; and in particular, functional MRI) became the dominant technique for investigating this organization non-invasively and *in vivo* (Eickhoff et al. 2018). Although regional delineations inferred based on architectonic criteria (e.g., cytoarchitecture) generally agree well with information gathered from MRI/fMRI (Amunts and Zilles 2015), such correspondences should be always interpreted with caution. These criteria may be weak predictors of functional organization in highly heterogeneous regions, for example when regions sit at the boundary of different Brodmann areas (BA; Brodmann 1909). Moreover, this relationship may be affected by strong inter-individual differences, which were not taken into account in most of the previous invasive studies characterized by relatively small sample sizes (Amunts and Zilles 2015). In addition to the previous prevalent invasive and lesion-based approaches, another way of conceptualizing functional organization and, more in general, the relationship between cognitive processes and their neural substrate, emerged from fMRI research with the functional localization approach (Kanwisher 2010). Specialized computations are performed by brain regions that can be reliably identified across individuals with fMRI using standard tasks (thus usually referred to as functional localizers; Kanwisher et al. 1997; Kanwisher 2010; O’Craven et al. 1999; Peelen and Downing 2005). In combination with the functional localization approach, research on structural MRI has shown that, despite the remarkable inter-individual variability in the organization of the gyri and sulci across the whole cortex (Desikan et al. 2006; Destrieux et al. 2010; Ono et al. 1990; Petrides 2018), these functional modules can also be localized based on anatomical landmarks (Fischl et al. 2008), which suggests a developmental link between the functional differentiation of brain regions and the mechanisms of cortical maturation (Zilles et al. 2013).

In sum, in the human brain, functional specialization appears to be tightly linked and possibly follows from brain structure, although it remains to be established exactly to what degree this principle holds within specific systems. In the prefrontal cortex (PFC), two structures, the frontal eye field (FEF) and the inferior frontal junction (IFJ) have largely overlapping but complementary roles, being involved in several orchestrating functions such as attention, working memory, cognitive control, and other top-down processes (Baldauf and Desimone 2014; Bedini and Baldauf 2021). In our previous review, we have in particular argued that despite these overlaps, it is their patterns of neural selectivity to spatial (FEF) vs non-spatial information (IFJ) that allows the dissociation of their role across these different functions (Bedini and Baldauf 2021). This functional specialization may in turn provide an effective way to localize these regions with fMRI. While the FEF has been studied extensively both in human and non-human primates, its precise localization and relationship to sulcal morphology in humans, and correspondence to the macaque FEF has proven to be difficult to establish (Amiez and Petrides, 2009; Petit and Pouget 2019; Schall et al. 2020; Tehovnik et al. 2000; but see Koyama et al. 2004). Evidence also suggests that FEF localization may be affected by substantial individual differences (Amiez et al. 2006; Kastner et al. 2007; Paus 1996; see Bedini and Baldauf 2021, for a discussion of this issue in the context of the multimodal parcellation (MMP1) by Glasser et al. (2016). The prevailing view is that the human FEF lies in the ventral bank of the superior precentral sulcus (sPCS), near its junction with the superior frontal sulcus (SFS; see Paus 1996 and Vernet et al. 2014, for a meta-analysis and a review of FEF localization, respectively). However, some authors have argued that instead, a region localized ventrally in the dorsal branch of the inferior PCS (iPCS), termed the inferior FEF (iFEF, or sometimes the lateral FEF) may be the putative homolog of the macaque FEF (Kastner et al. 2007; Schall et al. 2020). Moreover, it has been raised the related question of whether the inferior FEF has been under-reported in the fMRI literature (Derrfuss et al. 2012). In topographic mapping studies, peaks corresponding to the iFEF have been already reported (Kastner et al. 2007; Mackey et al. 2017) albeit they were not as consistent as FEF peaks in their presence across subjects and relative localization. Moreover, at least one study has previously reported activations in the iFEF using a saccadic localizer task, which were clearly segregated from those elicited by a Stroop task (Derrfuss et al. 2012). These analyses were performed in native space on an individual-subject basis, which is a very powerful approach that allows for carefully studying dissociations in adjacent neuroanatomical regions (Fedorenko 2021). The IFJ, a region found ventrally and anteriorly relative to the iFEF, is typically localized near the junction of the iPCS with the inferior frontal sulcus (IFS), sometimes encroaching into its caudal bank (Derrfuss et al. 2005). The IFJ was only much more recently characterized as a separate brain region (based on structural; Amunts et al. 2006; and functional criteria; Brass et al. 2005) that performs both specialized (Baldauf and Desimone, 2014; Bedini and Baldauf 2021) and general domain computations (Assem et al. 2020; Derrfuss et al. 2005), in line with the multiple demand hypothesis (Duncan 2010). Currently, however, due to the interspersed and close arrangement of specialized and multiple demand regions near the IFJ, common activation foci resulting from various cognitive processes have not been reported yet across experiments (see however Assem et al. 2020, for evidence from high-resolution fMRI data and improved inter-subject alignment methods).

Clearly, there is a need to better characterize the relationship between anatomy and functional specialization within the PFC. This is particularly critical, as the large inter-individual variability in the organization of the prefrontal areas and sulci (Germann et al. 2005; Juch et al. 2005) complicates the interpretation of the results of previous studies and may partly explain the discrepancies in the findings reported across research groups and methods. That such a link can in principle be successfully accomplished has been demonstrated in the visual system, where studies have shown that despite the inter-individual variability in the overall size, shape and position of the early visual cortex, specific anatomical landmarks (i.e., sulci) coincide very well with the borders of primary visual areas as derived from various sources of data, including cytoarchitecture, retinotopic mapping, myelin content and resting-state fMRI functional connectivity (Abdollahi et al. 2014; Fischl et al. 2008; Glasser et al. 2016; Sereno et al. 1995). For example, in Hinds et al. (2008) the authors used surface-based registration methods (Fischl et al. 1999) to identify V1 in new subjects from cortical folding information (i.e., the stria of Gennari) and showed that these methods outperformed volumetric methods in labeling this structure (see also Hinds et al. 2009). Similarly, Benson et al. (2012) used folding information to predict visual responses within the striate cortex to a retinotopic mapping fMRI protocol. When moving up into the cortical visual hierarchy, however, the relationships between cortical folding and other neuroanatomical information become more difficult to establish and interpret (Coalson et al. 2018; Glasser et al. 2016). Wang et al. (2015) created a probabilistic atlas of 25 topographic visual areas and showed that anatomical variability (as measured by the variance in gyral-sulcal convexity across subjects) and the overlap of functional activations (measured as peak probability values) were negatively correlated, particularly in higher-order visual areas, suggesting that the former may play an important role in shaping functional organization. Similarly, in one of the most comprehensive efforts to parcel the cortical surface with high-resolution non-invasive methods, Glasser et al. (2016) found that the lateral PFC is one of the brain districts where the intrinsic neuroanatomical variability is higher than in the rest of the brain (Glasser et al. 2016; Juch et al. 2005), as measured by a decrease in the test-retest reliability of their parcellation. While the former limitations (i.e., the weaker association between cortical folding and function, and inter-individual variability, which particularly affects volumetric group-level analyses; Coalson et al. 2018) have posed significant challenges to the interpretation of the relationship between cortical folding and functional specialization in higher-order visual regions, several studies have shown that adopting an individual-level approach in defining sulci may bear important implications for understanding cognitive function within the PFC (Amiez et al. 2006; Amiez and Petrides 2018; Derrfuss et al. 2009; Fedorenko 2021; Miller et al. 2021). For example, the study by Frost and Goebel (2012) showed that, by leveraging the former approach and improving the alignment in the cortical folding patterns (hence limiting the influence of the anatomical variability across subjects) using a technique termed curvature-driven cortex-based alignment, the overlap in FEF localization increased by 66.7% in the left hemisphere and 106.5% in the right hemisphere compared to volume-based registration in a sample of 10 subjects. These results suggest that the FEF is indeed strongly bound to a macro-anatomical location (i.e., the junction of the sPCS and the SFS; Paus 1996), and more generally the presence of a strong structure-to-function relationship in this region (see also Wang et al. 2015). In the study by Derrfuss et al. (2009), which again was carried out using an individual-level approach, 13 out of 14 subjects showed activations localized between the caudal bank of the IFS and the iPCS that corresponded to the anatomical description of the IFJ in a task-switching paradigm. Taken together, these studies point to the need to better characterize the relationship between sulcal morphology and functional specialization within the PFC. This research line may in the future allow predicting functional activity from neuroanatomical information alone, thus accomplishing one of the fundamental goals of contemporary cognitive neuroscience in terms of inferring structure-to-function relationships (Felleman and Van Essen 1991; Osher et al. 2016; Passingham et al. 2002; Saygin et al. 2012; Young et al. 2000). In summary, the organization of the regions localized along the banks of the major sulci of the posterior-lateral PFC (plPFC), namely the SFS, the sPCS, the iPCS, and the IFS, has yet to be clarified spatially.

In this study, we aimed to accurately localize the FEF and IFJ based on standard fMRI localizer tasks. In particular, we aimed to reassess the precise localization of the FEF in standard space, and its relationship with the localization of the iFEF as inferred using saccadic localizer tasks in the light of recent fMRI evidence (see Grosbras et al. 2005; Paus 1996 for previous meta-analyses using fMRI and PET experiments). Further, we also aimed at re-examining the precise localization of the IFJ in standard space by inferring the convergence of activations across several paradigms (see Derrfuss et al. 2005, for the convergence across task-switching and Stroop paradigms). This information may provide important clues on how to better interpret activations in the posterior-lateral PFC based on combined structural and functional criteria. Coordinate-based meta-analyses offer a convenient way to summarize and model the uncertainty in the activations found across several PET/fMRI experiments (Fox et al. 2014) based on specific paradigms and contrasts of interest, overcoming inter-individual variability, and allow to establish adequately powered brain-behavior relationship. The activation likelihood estimation (ALE) technique in particular allows inferring the spatial convergence of the foci reported in several independent fMRI experiments (Eickhoff et al. 2009, 2012). Here, we employed the ALE meta-analytic technique to accurately infer the localization of the FEF and IFJ activation peaks in standard space, thus aiming at resolving the discrepancies in the previous literature (in particular, concerning the precise localization of the FEF and IFJ). To do so, we analyzed data from standard functional localizers to better overcome the issue of inter-individual variability in the localization of these regions. These are well-validated tasks specifically designed to target pre-defined ROIs (Fedorenko 2021; Kanwisher 2010) and are typically performed independently from the main fMRI task. By using the inferred peak coordinates as seeds, we also performed a meta-analytic connectivity modeling (MACM) analysis (Robinson et al. 2010) to investigate the coactivation profiles of FEF and IFJ in fMRI studies across paradigms. Overall, our study aims to provide meta-analytic groundwork for investigating the relationship between functional specialization and connectivity of the FEF and IFJ in large neuroimaging datasets (e.g., the Human Connectome Project; Van Essen et al. 2013). Additionally, it also aims at offering some consensus in the field and anatomical priors to accurately localize these regions with fMRI and to guide future non-invasive brain stimulation studies.

## Materials and methods

### Activation likelihood estimation fMRI meta-analysis method

The ALE is a powerful meta-analytic technique that allows for assessing the spatial convergence of the activations reported in the neuroimaging literature (Eickhoff et al. 2009, 2012). As a coordinate-based technique, ALE takes as input the activation peaks reported by several independent neuroimaging studies and tests their significance against a null distribution of the foci across the whole brain (Eickhoff et al. 2012). This ALE feature is particularly useful given that in the neuroimaging literature results are usually reported and summarized as x, y, z coordinates in standard space (Talairach or MNI), rather than as full activation maps accompanied by a statistical summary of the effect sizes, and even more rarely shared in that form (for important initiatives in neuroimaging data sharing see however NeuroVault: https://neurovault.org/, Gorgolewski et al. 2015; OpenNeuro: https://openneuro.org/, Markiewicz et al. 2021; BALSA: https://balsa.wustl.edu/, Van Essen et al. 2017, and Anima: https://anima.inm7.de/index.php, Reid et al. 2015, among other initiatives). This aspect becomes crucial in the case of brain regions that may be under-reported in the fMRI literature (such as the iFEF; Derrfuss et al. 2012; Kastner et al. 2007) or which only recently began to be included in the brain atlases taxonomy (such as the IFJ; Bedini and Baldauf 2021; Sundermann and Pfleiderer 2012). Here, we exploited this ALE feature by applying this technique to analyze two independent collections of fMRI studies performed over the last 30 years with the primary aim of accurately inferring FEF and IFJ localization in MNI152 space using the GingerALE software (v. 3.0.2; https://www.brainmap.org/ale/). In the ALE procedure, each set of foci reported in a study is modeled as a three-dimensional Gaussian distribution centered around the coordinates and whose size is determined based on the experiment sample size (Eickhoff et al. 2012). In particular, larger sample sizes result in tighter Gaussians, which reflects lower uncertainty about the ‘true’ location reported, whereas lower samples lead to larger Gaussians that are more spread around the respective peak coordinates, thus conveniently reflecting lower confidence about their corresponding locations. These activations are then combined into a modeled activation map for each experiment of a study. Importantly, in the revised ALE algorithm, within-study effects that could result from the summation of adjacent foci are minimized, so that studies that reported activation in a higher number or more densely organized foci won’t drive the ALE results disproportionately (Turkeltaub et al. 2012). By computing the union of all these modeled activation maps, an ALE score for each voxel in the brain is obtained (Eickhoff et al. 2009, 2012). The significance of these scores is then assessed by comparing them with the null distribution obtained by randomly reassigning the modeled activations across the whole brain with a permutation approach. Finally, the thresholded p-values are usually corrected for multiple comparisons using either voxel-level or cluster-level family-wise error (FWE). The use of uncorrected p-values and false discovery rate is instead generally not advised since it can lead to spurious findings (Eickhoff et al. 2016).

The present meta-analysis focused on specific cognitive functions (described more in detail in the ‘Study selection criteria’) in which the FEF and IFJ and the associated brain networks are relatively well-known to be involved. More specifically, in what we will refer to in the following as the ‘FEF sample’, we applied the ALE technique to several independent fMRI studies requiring the planning and execution of visually-guided and voluntary eye movements, as a considerable number of previous studies clearly showed that these types of tasks elicit activation in the FEF, and other eye fields (for previous meta-analyses see Cieslik et al. 2016; Grosbras et al. 2005; Jamadar et al. 2013; Paus 1996). In particular, tasks requiring the execution of prosaccades and antisaccades contrasted against a fixation baseline are the most prevalent and consensually established FEF functional localizer in the human fMRI literature (Amiez et al. 2006; Amiez and Petrides 2018; Kastner et al. 2007). In the case of the ‘IFJ sample’, it is arguably more difficult to pinpoint a universally accepted functional localizer for this region, and in fact, very few studies have implemented a separate and targeted localizer paradigm to isolate the IFJ (for relevant examples see Baldauf and Desimone 2014; Zanto et al. 2010; Zhang et al. 2018) for subsequent analyses. In addition, there is increasing evidence that the IFJ may be characterized as a region more generally involved in different and often overlapping functions (i.e., a multiple demand region; Assem et al. 2020; Derrfuss et al. 2005; Sundermann and Pfleiderer 2012), which may even further complicate the definition of a standard localization method. Therefore, in the ‘IFJ sample,’ we anticipate that we analyzed data from a more heterogeneous collection of fMRI studies investigating covert attention, working memory, and cognitive control across a wider range of paradigms. Whenever the sample sizes of the studies retrieved allowed for it, we also carried out post hoc control analyses to examine potential spatial discrepancies between the IFJ peaks derived from splitting up the full localizer sample according to specific cognitive functions and paradigms (see Supplementary Information p. 5 and 14-15).

### Study selection criteria

The selection criteria of the sample of studies for the present meta-analysis followed the best-practice recommendations and guidelines by Müller et al. (2018). Multiple bibliographic searches were performed between May 2019 and January 2021 (cutoff date). A final search was conducted with the same criteria and cutoff dates (i.e., 1st of January 1990 - 1st of January 2021) by the first author MB to comply with the updated PRISMA guidelines (Page et al. 2021) and as a sanity check. The selection procedure is reported in Figures S1 and S2 (created based on the PRISMA flow diagram; Page et al. 2021), which refer to the ‘FEF sample’ and the ‘IFJ sample’, respectively. All the bibliographical searches were carried out using Web of Science (https://www.webofscience.com). We searched records in the Web of Science Core Collection using the keywords ‘fMRI’ AND ‘frontal eye field’ (all fields) in the first instance, and ‘fMRI’ AND ‘inferior frontal junction’ (all fields) in the second. We complemented these results with other sources (Google Scholar, personal collection of articles and references cited by the studies retrieved) by one of the authors (MB). In the FEF sample, our search identified a total of 711 records, from which we removed all the review papers. 665 records were further screened, and 470 of these were sought for retrieval to assess their adequacy with respect to the inclusion criteria (described below). In the IFJ sample, 375 results were identified, from which, after removing review papers, 356 records were further screened, and 142 of these were sought for retrieval to assess their adequacy with respect to our original inclusion criteria. The general inclusion criteria consisted of the following. Each study selected: 1. Reported coordinates in standard space (either MNI or Talairach); 2. Was an fMRI study (we decided to not include positron emission tomography studies as our goal was to keep our samples as homogeneous as possible in terms of the signal measured, the spatial resolution and analysis pipelines (Botvinik-Nezer et al. 2020) to make our results specific to the fMRI field); 3. Performed on a scanner of 3T or higher field; 4. Tested and reported results from healthy adults (18-60 years old; or an appropriate control group in the case of clinical studies); 5. The study acquired whole-brain fMRI data or with a FOV that was sufficiently large to cover the frontal cortex and in particular its posterior aspect (see the Supplementary Tables 1 and 2 for the FOV of each study). Even though the latter criterion would lead to the inclusion of studies with partial brain coverage, which is generally not recommended in ALE analyses (Müller et al. 2018), given that our main research question focused on the standard localization of the FEF and IFJ, we motivate it by accepting the tradeoff derived from having access to a larger sample for these regions, as opposed to having less sensitivity in detecting other regions that are consistently active during the tasks included (which however are not the main focus of the present study), but not reported simply due to the lack of brain coverage. In the Supplementary Information, we report the results of a control analysis excluding first studies with partial brain coverage and second, studies performing ROI analyses, as per the best practices recommended by Müller et al. (2018). The last group of inclusion criteria is specific to each sample (the FEF or the IFJ) and is primarily related to the type of experimental paradigm utilized in the fMRI study and the specific contrasts analyzed (described more in detail below). Here, we strived to find a balance that would adequately represent the various localization methods that have been pursued in the fMRI literature, while also assigning a higher weight in the sample to the more standardized and replicated localization approaches.

### FEF sample inclusion criteria

The human FEF is a well-characterized region in the fMRI literature (Bedini and Baldauf 2021), although some uncertainties persist regarding the correspondence of its localization obtained from fMRI compared with other methods (i.e., brain stimulation; Vernet et al. 2014) and with the macaque FEF (Koyama et al. 2004; Petit and Pouget 2019). The region is crucially involved in the top-down control of eye movements and spatial attention (Astafiev et al. 2003; Beauchamp et al. 2001; Corbetta et al. 1998), and it is considered a prominent node of the dorsal attention network (Corbetta and Shulman 2002; De Pasquale et al. 2010; Fox et al. 2006; Yeo et al. 2011). A very simple and time-efficient yet effective way to localize the FEF in fMRI scans is to have participants perform an experimental block of visually-guided saccades towards an unpredictable peripheral target, and contrast this activation with a fixation block (Amiez and Petrides 2006). The resulting activations - usually found near the junction of the SFS and the sPCS - are then assumed to correspond to the FEF (Paus 1996). However, depending on the statistical thresholds and analytical approach adopted, in addition to this superior cluster, often this type of contrast reveals a more widespread pattern of activity along the banks of the iPCS (Beauchamp et al. 2001; Kastner et al. 2007). Therefore, this localization method doesn’t seem to have adequate functional specificity if not combined with the additional anatomical criteria mentioned above. Building on this approach, the antisaccade task and its neural mechanisms have been extensively studied in the primate neurophysiology literature (Munoz and Everling 2004), and this task has been employed as a measure of inhibitory control in healthy and clinical populations in humans (Hutton and Ettinger 2006). Briefly, in the antisaccade task, the subject is required to keep fixation until a visual target appears and to look at its mirror location (Hallet 1978). Computationally, this requires at least two mechanisms: the first one inhibits a reflexive saccade towards the visual onset, and the second is responsible for executing a saccade towards the opposite location (the endpoint is in this case endogenously generated; Munoz and Everling 2004). An interesting feature of this task is that compared to the prosaccade task, it gives rise to frequent direction errors, which reflect a failure to inhibit reflexive behavior (Hutton and Ettinger 2006; Munoz and Everling 2004; Pierrot-Deselleigny et al. 2002). Inserting a short (e.g., 200 ms) gap period before the presentation of the target decreases saccadic reaction times (Saslow 1967) and further increases the ratio of these direction errors (Munoz and Everling 2004). fMRI studies comparing the regions involved in prosaccade vs antisaccade task performance have found overlapping activations in the FEF in both tasks, although the antisaccade task recruits additional regions that seem to reflect the increased executive demands of this task (McDowell et al. 2008). Within FEF, there is also increased activity in the antisaccade compared to the prosaccade task (Connolly et al. 2000). This is particularly evident during the preparatory phase (Brown et al. 2006; Connolly et al. 2002; Curtis and Connolly 2008; DeSouza et al. 2003: Ford et al. 2005) and when the two tasks are presented in a mixed fashion (Pierce and McDowell 2015, 2017). Based on these results, it could be hypothesized that contrasting antisaccades vs prosaccades may offer better specificity to localize clusters of activity within the FEF compared to the prosaccade vs fixation blocked design described earlier. Previous studies didn’t address this hypothesis directly, and it still needs to be tested in a within-subject design by comparing the activation topographies at the individual subject level. Finally, in modified versions of the spatial cueing paradigm (Fan et al. 2005; Posner 1980), univariate analyses contrasting valid vs neutral/invalid trials are often used to localize all the main regions belonging to the dorsal attention network (Corbetta and Shulman 2002), which are subsequently used as ROIs for functional and effective connectivity analyses (for example, see Vossel et al. 2012, and Wen et al. 2012). It can be argued that, even though these adaptations are not generally employed as independent functional localizers for the FEF, they may be well adept to isolate this region under the assumption that covert and overt shifts of spatial attention have a shared and overlapping source in this region, which seems well supported by fMRI (Astafiev et al. 2003; Beauchamp et al. 2001; Corbetta et al. 1998; Jerde et al. 2012) and comparative evidence (Buschman and Miller 2009; Moore and Fallah 2001; reviewed in Fiebelkorn and Kastner 2020). Indeed, the studies that directly investigated this question generally reported a strong degree of spatial overlap, although they also suggest that the signal measured in covert paradigms tends to be weaker than in overt tasks (Beauchamp et al. 2001; De Haan et al. 2008) and thus possibly less robust across fMRI data analysis pipelines (Botvinik-Nezer et al. 2020). Thus, an open question is whether oculomotor and covert spatial attention tasks are equally efficient in localizing the FEF.

In summary, for the reasons introduced above, we included in the FEF sample all the studies that investigated the planning and execution of visually-guided and voluntary eye movements (prosaccades and antisaccades) as well as covert spatial attention using both blocked and event-related designs, analyzing mainly the following contrasts: 1. prosaccades > fixation; 2. antisaccades > fixation; 3. prosaccades & antisaccades > fixation; 4. anti-saccades > pro-saccades; 5. valid > neutral/invalid trials (see Figure S1 for an overview of the selection procedure following the PRISMA2020 guidelines; Page et al. 2021; for the full list of studies and the related information see Tables S1 and S2 in the Supplementary Information). Combining these contrasts allowed us to carry out our main analysis complemented by three control analyses, respectively designed to replicate a previous study (Cieslik et al. 2016) and to investigate two additional research questions (see Supplementary Information p. 4). In our main localizer analysis, we pooled together all studies that reported at least a contrast related to the planning and execution of prosaccades, both prosaccades and antisaccades, and antisaccades, contrasted with a fixation baseline.

### IFJ sample inclusion criteria

In contrast to the FEF, the IFJ doesn’t have a broadly accepted homolog in the macaque (Bedini and Baldauf 2021; see however Bichot et al. 2015, 2019; and Neubert et al. 2014) and its role started to be investigated only much more recently with fMRI (Brass et al. 2005). Its functional profile remains to date not well understood and is characterized by a remarkable functional heterogeneity (Muhle-Karbe et al. 2016; Ngo et al. 2019). Consistent with this idea, recent high-resolution fMRI studies showed that the IFJ (and in particular, the posterior IFJ as defined according to the MMP1 by Glasser et al. 2016) belongs to the core multiple-demand system of the brain (Assem et al. 2020, 2021), which identifies a set of regions that are engaged in multiple processes often across different cognitive domains (Duncan 2010). This particular position in the cognitive processing architecture arguably poses a severe challenge in trying to define a gold standard for an fMRI localization method for this region, which would allow for effectively segregating it from adjacent coactive regions (for an excellent example of such an approach see however Derrfuss et al. 2012; we return to this point in the discussion). Several promising approaches to localize the IFJ both at the individual and at the group level have nevertheless previously been reported from different research groups ranging from attention and working memory to cognitive control paradigms. For example, the series of studies from the group led by Brass and colleagues were critical in establishing the IFJ as a region involved in task preparation and more generally in cognitive control (Brass and Von Cramon, 2002, 2004; Derrfuss et al. 2004; see also Cole and Schneider 2007).

In summary, based on the evidence discussed above, in the IFJ sample, we decided to include attentional (i.e., rapid serial visual presentation (RSVP) and endogenous cueing paradigms), working memory (primarily n-back paradigms), and cognitive control paradigms (i.e., task-switching and Stroop tasks). These inclusion criteria were based on Derrfuss et al. (2005), who investigated switching and Stroop paradigms, and we extended them to attentional and working memory paradigms that presumably also tap cognitive control functions. These paradigms all stress the maintenance of a task set to control behavior, which should be one of the integrative IFJ functions according to our hypothesis. The main contrasts analyzed were therefore quite heterogeneous, but can be broadly grouped into the following primary ones (see Figure S2 for an overview of the selection procedure following the PRISMA2020 guidelines; Page et al. 2021; for the full list of studies and the related information see Table S2): 1. Oddball > Target trials in covert attention paradigms (e.g., RSVP paradigms); 2. Functional connectivity with a seed perceptual region (e.g., V4, V5, FFA) in the Attend > Ignore condition in n-back paradigms; 3. Switching > Repetition trials in task-switching paradigms; 4. Incongruent > Congruent trials in Stroop paradigms (see the Supplementary Information for control and contrast analyses).

In conclusion, the final sample of the included papers for our ALE meta-analysis was n = 51 for the FEF, and n = 30 for the IFJ sample (see the Supplementary Information, Figures S1 and S2, and Tables S1 and S2 for a summary of the studies). The number of experiments was 35 for the FEF localizer sample, and 32 for the IFJ localizer sample. Both sample sizes were within the recommended range (i.e., a minimum of 17-20 experiments) to have adequate statistical power with ALE as derived from empirical simulations (Eickhoff et al. 2016).

### Activation likelihood estimation procedure

We extracted all the foci from the studies included in the FEF and IFJ samples, and we converted all the Talairach coordinates to the MNI152 space using the Lancaster transform as implemented by the function provided in the GingerALE software (v. 3.0.2; Laird et al. 2010; Lancaster et al. 2007). We note that after the SPM2 version, the MNI templates distributed are consistent across FSL and SPM software packages, being compliant with the ICBM-152 coordinate space (Fonov et al. 2009), so for any later version of these packages we used the Talairach to MNI FSL transform. Where other software packages were used for spatial normalization, we again employed the Talairach to MNI FSL transform for consistency, as the other transformation provided in GingerALE represents a pooled FSL/SPM transformation (Lancaster et al. 2007) that would only lead to systematic displacement of the coordinates. In only two cases (i.e., Manoach et al. 2007; Mao et al. 2007) the studies employed the mapping from MNI to Talairach developed by Brett et al. (2002). These coordinates were therefore mapped back to the MNI space using this specific transformation, as recommended in the GingerALE user manual.

For the main localizer analyses (FEF and IFJ localizer samples), the ALE parameters were set to 10000 threshold permutations (as recommended in Eickhoff et al. 2016) and a voxel-level FWE of 0.01 was applied (Eickhoff et al. 2017) with a minimum cluster size of 50mm^3^ (corresponding to 6 voxels). Compared to cluster-level FWE inference, which can only allow inferring that a given cluster is above a significance threshold as a whole, but critically, not that any putative region that is included in the cluster is individually significant on its own (Eickhoff et al. 2016), voxel-level FWE allows to more readily interpret all the cluster extent as well as its peak location from the main localizer samples anatomically (Eickhoff et al. 2016). Moreover, our sample sizes ensured that these clusters would not be driven by a contribution exceeding 50% of any individual study. Therefore we would like to stress that performing the ALE procedure for the main localizer samples using voxel-level FWE arguably represents the most conservative approach to inferring spatial convergence in our samples and allows us to interpret individual voxel ALE values as a proxy for the most active location across experiments. When retrieving the relevant foci, we first grouped the studies by subject group rather than by experiment (Turkeltaub et al. 2012). This was done because grouping by subject group further minimizes within-study effects (Turkeltaub et al. 2012). When a single experiment reported multiple contrasts of interest, we therefore pooled them under the same subject group. We note that however, in all cases in which the studies reported more than one contrast of interest they were drawn from the same experiment (with very few exceptions; see Tables S1 and S2), so our strategy didn’t unfairly pool together partially independent observations and was practically almost equivalent to grouping by experiment. When an experiment failed to report significant activation for some ROIs, we used the lower number of subjects that had above threshold activations in all ROIs from a contrast of interest if this information was available. In summary, our study grouping strategy allowed us to fully exploit the information gathered from the different contrasts that were performed in the original studies and also to carry out our control analyses with the largest possible sample sizes. To validate the results of the main ALE analyses and to further assess the reliability of the ALE peaks found, we also carried out a leave-one-experiment-out procedure (LOEO; Eickhoff et al. 2016) on the main FEF and IFJ localizer samples using the same foci grouping strategy. Since we found identical ALE peaks as in the main ALE analyses using 1000 threshold permutations, we performed the LOEO procedure with the same parameter to reduce computational times.

### Comparison method of the ALE clusters and peaks with previous coordinate-based meta-analyses, relationship to macro-anatomical information and the MMP1

To interpret our results more carefully, we compared the significant activation clusters from our ALE main localizer analyses with the results from previous meta-analyses results and brain atlases (Derrfuss et al. 2005; Glasser et al. 2016; Klein et al. 2012; Paus 1996). First, we described the anatomical location of each cluster and assigned the corresponding Brodmann label using the Talairach Daemon in GingerALE (Lancaster et al. 2000). Second, to compare our results with previous meta-analyses (Derrfuss et al. 2005; Paus 1996), we mapped our ALE peaks to the Talairach space using the transformation developed by Lancaster et al. (2007) as implemented in GingerALE (MNI (FSL) to Talairach; see Table 3). Third, to relate our results to surface-based atlases (Klein et al. 2012; Glasser et al. 2016), we followed two distinct approaches. The Mindboggle 101 atlas (Klein et al. 2012) describes the macro-anatomical organization of the human brain as delineated by sulcal and gyral information. The atlas was recently mapped to the MNI152 non-linear symmetric template (Manera et al. 2020), so we manually imported this atlas in FSL and in FSLeyes as described in this GitHub repository: The-Mindboggle-101-atlas-in-FSL. We assigned one of the Mindboggle 101 labels to each of the ALE peak coordinates using the atlasquery command-line tool with FSL (v. 6.0.3; Jenkinson et al. 2012). For atlases that were released and best interpreted in a surface-based format, such as the MMP1 (Glasser et al. 2016; see Coalson et al. 2018 for an in-depth discussion), we instead employed the mapping technique developed in Wu et al. (2017) to register our ALE results from the MNI152 space (Fonov et al. 2009) to FSaverage (Fischl et al. 1999). A version of the MMP1 (Glasser et al. 2016) mapped to the FSaverage surface was made available using the method described in Mills (2016; https://figshare.com/articles/dataset/HCP-MMP1_0_projected_on_fsaverage/3498446). Once we mapped the ALE clusters to this surface, we also mapped the MNI152 coordinates corresponding to each ALE peak to a vertex on the inflated surface (Wu et al. 2017) and we assigned each of these to the respective MMP1 labels (Table 5). To describe the anatomical labels associated with each ALE cluster using more specific labels (compared to the Talairach Daemon), we used a volumetric version of this atlas for convenience. The source files that were used to import the atlas are the same as in Huang et al. (2021). The volumetric version of the MMP1 was manually imported in FSLeyes as described in this GitHub repository: The-HCP-MMP1.0-atlas-in-FSL).

### Meta-analytic connectivity modeling method

After obtaining the ALE peaks from the main localizer analyses, we exploited this information to perform a data-driven analysis of the coactivation patterns of the FEF and the IFJ across the whole brain to uncover their task-based fMRI functional connectivity fingerprint (Langner and Camilleri 2021; Passingham and Wise 2002). We therefore retrieved all the papers matching specific criteria (described below) from the BrainMap database using Sleuth (https://www.brainmap.org/sleuth/; Fox and Lancaster 2002), and we analyzed these foci by employing the MACM technique (Robinson et al. 2010). This technique leverages the ALE algorithm and allows inferring all the regions that coactivate with a given seed region that is selected a priori. This analysis also allowed us to perform a reverse inference on these coactivation patterns (Poldrack 2011). More specifically, we sought to functionally decode and characterize the various behavioral domains that are associated above chance with each of these by using a standardized taxonomy (Fox et al. 2005) via the Mango software (v. 4.1) behavioral analysis plugin (v. 3.1; Lancaster et al. 2012).

The studies were retrieved from the BrainMap database using Sleuth according to the following fields (all linked using the ‘AND’ operator) and specifications: in the Experiment field, the “context” field was set to “normal mapping”, in the “activation” field we searched for “activations only”, with “Imaging modality” being set to “fMRI”. Finally, four separate searches were conducted in the BrainMap database by setting the “locations” field as corresponding to each left hemisphere (LH) and right hemisphere (RH) seed region (LH FEF, RH FEF, LH IFJ, and RH IFJ). We first transformed each seed location from MNI152 to Talairach space (which is the standard in Sleuth and also used internally by Mango’s behavioral plugin) using the transformation by Lancaster et al. (2007; the FSL transformation) and we created cuboid seeds of 6 mm centered around the respective ALE peaks. With these criteria, we were able to retrieve a range of 19 to 53 studies across seed locations. In particular, we retrieved 26 studies (27 experiments) for the LH FEF seed, 19 studies (19 experiments) for the RH FEF seed, 53 studies (59 experiments) for the LH IFJ, and 31 studies (31 experiments) for the RH IFJ. We note that the different number of studies retrieved possibly reflects a combination of the increased base probability of finding activations within a specific ROI (Langner et al. 2014) but also the fact that some ROIs tend to participate in multiple functional networks (Langner and Camilleri 2021). Thus, while our sample size was generally greater in the case of the IFJ, this information matches what was previously reported about the coactivation patterns of FEF and IFJ (the IFJ being part of the frontoparietal network, that has been previously shown to work as a hub network; Cole et al. 2013). More importantly, these sample sizes allowed for adequately powered inference using ALE (Eickhoff et al. 2016). Therefore, we used all the foci retrieved separately from each seed location as inputs for GingerALE. The ALE parameters were set to an 0.001 uncorrected p-value, 1000 threshold permutations and a cluster-level FWE of p < 0.01. In the functional decoding analysis, we used the same 6 mm cuboid seeds as in the MACM analysis centered around the respective FEF and IFJ ALE peaks. Associations with behavioral domains were considered statistically significant when their z-score was ≥ 3, corresponding to a threshold of p < 0.05 (Bonferroni corrected).

### Data availability

The results of this study will be made available in NeuroVault (https://neurovault.org/) upon article acceptance.

## Results

### FEF and IFJ localizer samples ALE main clusters

In the FEF localizer sample, the activations converged most strongly in two main clusters localized in the left and right posterior dorsolateral PFC. Two ALE peaks were found near the junction of the sPCS with the SFS, localized in the anterior (in the LH) and posterior (in the RH) banks of the sPCS (Figure 1). These peaks match well the classical description of the human FEF as inferred with fMRI (Petit and Pouget 2019; Vernet et al. 2014). Our LOEO procedure overall confirms the reliability of the localization of these ALE peaks (see Table 3; LH: 26/35; RH: 23/35). In the IFJ localizer sample, the activations converged more strongly in two main clusters localized in the left and right posterior ventrolateral PFC. These clusters extended both in the dorsal and ventral portion of the iPCS, partially encroaching on the IFS (see Figure 1). The cluster in the right hemisphere was slightly more focused spatially compared to the cluster in the left hemisphere. Crucially, in both clusters, we found that the ALE peaks were localized along the posterior bank of the iPCS, near its ventral junction with the IFS, which closely matches the description of the IFJ (Derrfuss et al. 2005; Muhle-Karbe et al. 2016). Again, our LOEO procedure overall suggests that these ALE peaks are highly reliable across experiments (see Table 3; LH: 31/32; RH: 24/32).

**Fig. 1.**
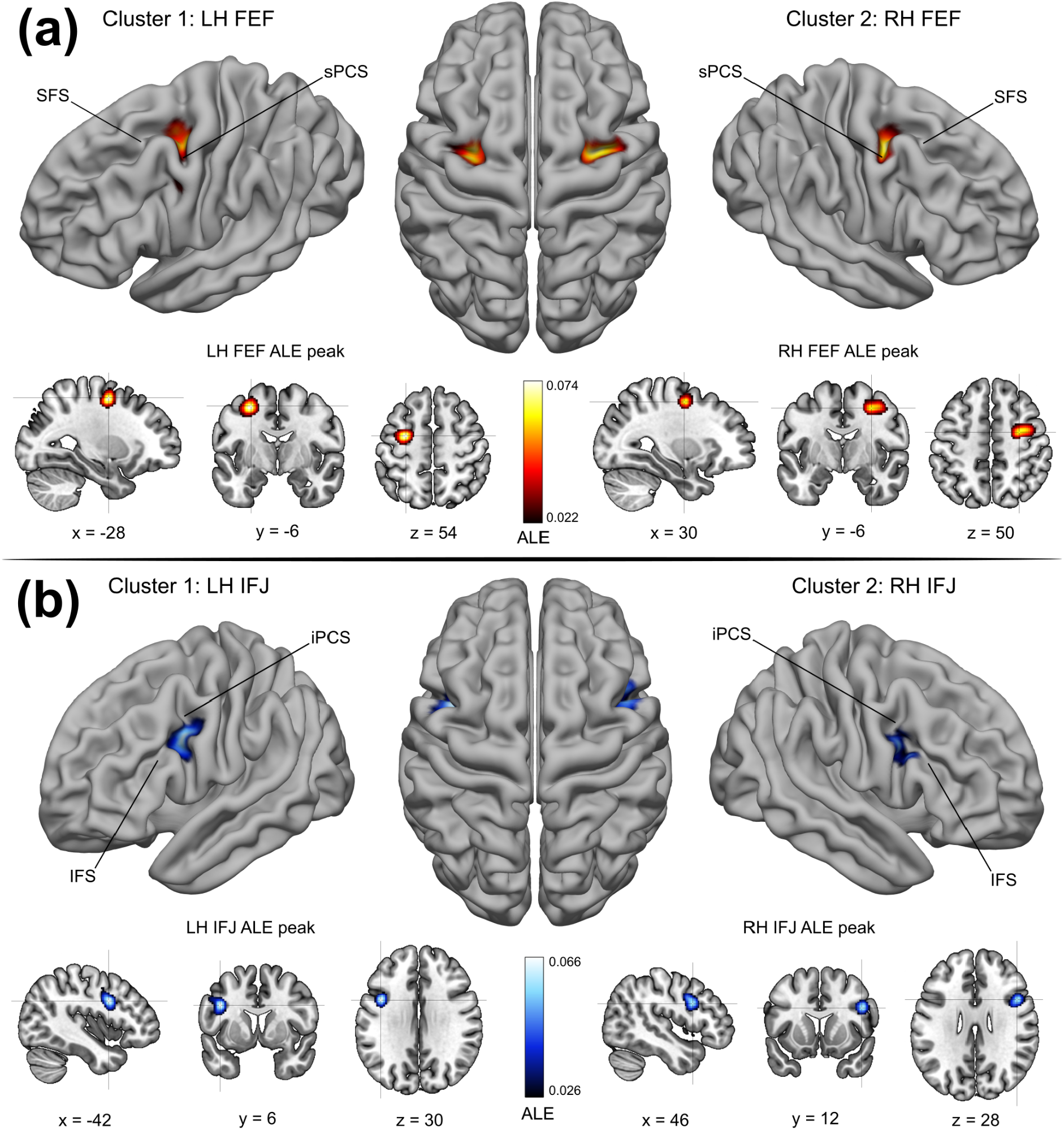
FEF and IFJ localizer samples - ALE main clusters. **A** ALE results from the FEF localizer sample. Two main clusters were found in the posterior dorsolateral PFC, which corresponds to the description of the anatomical location of the FEF (Paus 1996; Vernet et al. 2014). The FEF peaks were localized at the junction of the sPCS and the SFS, in the anterior (in the LH) and posterior (in the RH) banks of the sPCS. **B** ALE results from the IFJ localizer sample. Two main clusters were found in the posterior ventrolateral PFC, and their respective peaks were localized along the posterior bank of the iPCS, near its ventral junction with the IFS. The location of these peaks and the corresponding MNI coordinates match the description of the IFJ (Derrfuss et al. 2005; Muhle-Karbe et al. 2016)

### FEF localizer sample ALE results - FEF lateral peak and other significant clusters

In the left hemisphere, we also found a lateral peak within the main FEF cluster, which was localized on the bank of the iPCS, dorsal to its junction with the IFS (see Figure 2). This lateral peak corresponds to what has been previously referred to as the inferior or the lateral FEF (Derrfuss et al. 2012; Kastner et al. 2007; Luna et al. 1998). In addition to the main FEF clusters in the left/right PFC, the ALE technique revealed three other consistently activated clusters. These clusters were localized in the medial frontal gyrus and the left/right posterior parietal cortex (Table 1). The cluster in the medial frontal gyrus comprised the SCEF (Amiez and Petrides 2009) and the dorsal cingulate motor cortex (Glasser et al. 2016). In the posterior parietal cortex, two bilateral superior clusters spanned the precuneus and the SPL (Scheperjans et al. 2008a, 2008b), and an additional cluster was found in the right anterior intraparietal area (Glasser et al. 2016; Numssen et al. 2021).

**Fig. 2.**
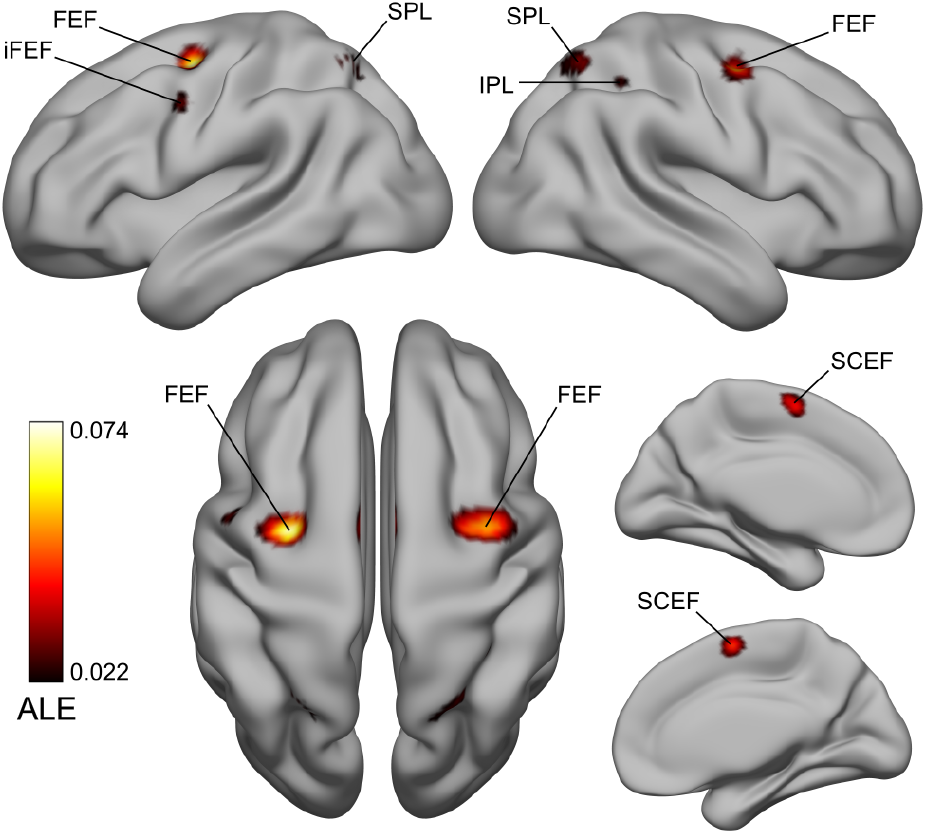
FEF localizer sample ALE results - FEF lateral peak and other significant clusters. In the FEF localizer analysis, we also found a lateral peak in the LH, which was localized near the bank of the iPCS, dorsal to its junction with the IFS, corresponding to the iFEF (Kastner et al. 2007; Luna et al. 1998). It is unclear whether this region should be considered part of the FEF proper (Glasser et al. 2016; Mackey et al. 2017). We found three other significant clusters localized in the supplementary and cingulate eye field (SCEF) and the dorsal cingulate motor cortex, and the precuneus/superior parietal lobule (SPL) and the right anterior intraparietal area

**Table 1.** FEF localizer sample ALE results

| Cluster | Macroanatomical location | Hemi | MNI152 coordinates |  |  | ALE value | Volume (mm<sup>3</sup>) | BA |
| --- | --- | --- | --- | --- | --- | --- | --- | --- |
|  |  |  | x | y | z |  |  |  |
| 1 | Precentral Gyrus | L | -28 | -6 | 54 | 0.0736 | 4784 | 6 |
|  | Precentral Gyrus | L | -48 | -2 | 40 | 0.0293 |  | 6 |
| 2 | Precentral Gyrus | R | 30 | -6 | 50 | 0.0623 | 4208 | 6 |
| 3 | Medial Frontal Gyrus | L/R | 0 | 0 | 58 | 0.0547 | 3016 | 6 |
| 4 | Precuneus | L | -22 | -58 | 56 | 0.0418 | 1512 | 7 |
| 5 | Precuneus | R | 24 | -60 | 56 | 0.0376 | 1056 | 7 |
| 6 | Inferior Parietal Lobule | R | 36 | -46 | 48 | 0.0262 | 168 | 40 |

### IFJ localizer sample ALE results - Other significant clusters

In addition to the main IFJ clusters in the left/right PFC, we found seven consistently activated clusters forming a broad fronto-parietal network (see Figure 3). In the frontal cortex, the first cluster was localized in the in the dorsal anterior cingulate cortex (dACC; Cole and Schneider 2007) and SCEF (Amiez and Petrides 2009), a second in the left precentral gyrus (within the putative FEF), and finally, two other clusters were localized in the bilateral insular cortex and claustrum (Table 2). Posteriorly, we also found a cluster in the left SPL/inferior parietal lobule (IPL), and a smaller cluster in the right SPL/IPL (Numssen et al. 2021; Scheperjans et al. 2008a, 2008b).

**Fig. 3.**
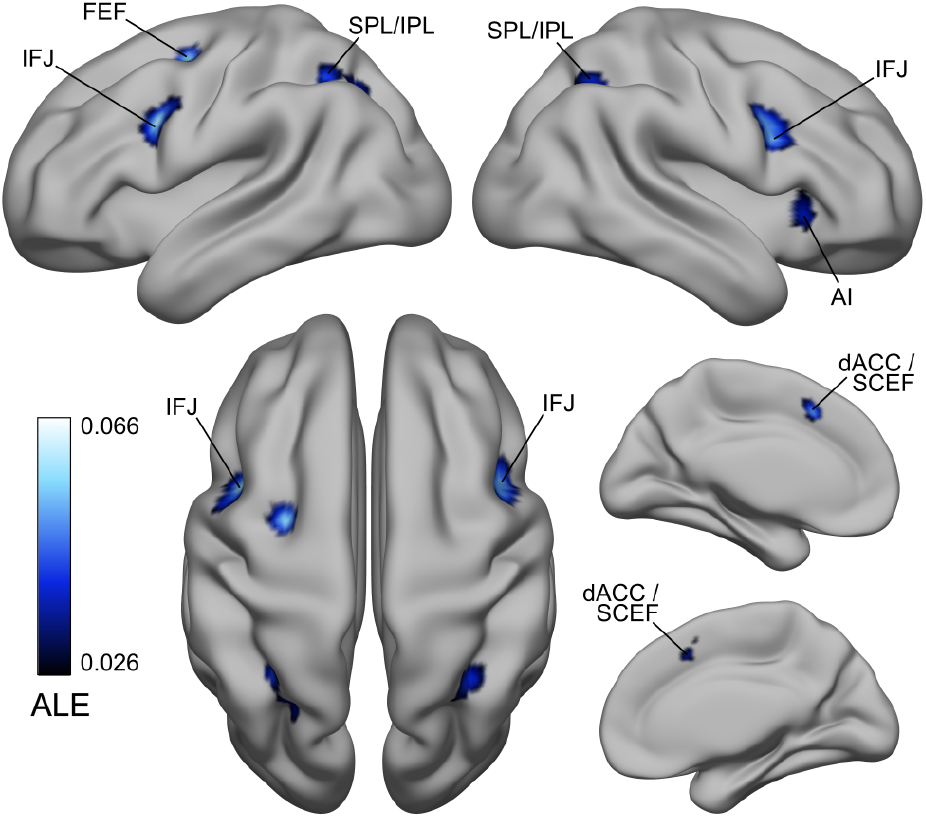
IFJ localizer sample ALE results - Other significant clusters. In addition to the bilateral IFJ clusters, we found significant activations in the dACC/SCEF, the left FEF, in two clusters in the insular cortex and claustrum (not visible in the left hemisphere), and finally, in the SPL/IPL. Given that these areas were activated across different paradigms, we suggest that they could be associated with the “encoding and updating of task-relevant representations” as first hypothesized by Derrfuss et al. (2005). From a network perspective, they belong to the frontoparietal network (Cole et al. 2013; Yeo et al. 2011)

**Table 2.** IFJ localizer sample ALE results

| Cluster | Macroanatomical location | Hemi | MNI152 coordinates |  |  | ALE value | Volume (mm<sup>3</sup>) | BA |
| --- | --- | --- | --- | --- | --- | --- | --- | --- |
|  |  |  | x | y | z |  |  |  |
| 1 | Precentral Gyrus | L | -42 | 6 | 30 | 0.0657 | 3488 | 6 |
| 2 | Inferior Frontal Gyrus | R | 46 | 12 | 28 | 0.0597 | 3016 | 9 |
| 3 | Medial Frontal Gyrus | L | -2 | 18 | 44 | 0.0573 | 2328 | 6 |
| 4 | Superior Parietal Lobule | L | -30 | -54 | 48 | 0.0444 | 1504 | 7 |
|  | Precuneus | L | -26 | -66 | 44 | 0.0388 |  | 7 |
| 5 | Precentral Gyrus | L | -28 | -4 | 54 | 0.0577 | 1208 | 6 |
| 6 | Superior Parietal Lobule | R | 34 | -56 | 48 | 0.0408 | 800 | 7 |
| 7 | Clastrum | R | 32 | 22 | -2 | 0.0385 | 728 | * |
| 8 | Clastrum | L | -30 | 18 | 2 | 0.0342 | 424 | * |

### Spatial relationship of the main FEF and IFJ clusters and ALE peaks with previous coordinate-based meta-analyses, macro-anatomical information and the MMP1

The comparison of the FEF and IFJ ALE peaks from the localizer samples analyses overall shows good spatial correspondence with results from previous meta-analyses and the MMP1 (Glasser et al. 2016), but with some important differences that are worth examining in detail (see Table 3). The FEF ALE peaks from our results are localized much more medially and posteriorly relative to the results reported in Paus (1996), highlighting marked spatial differences with this landmark FEF meta-analysis. In contrast, the IFJ ALE peaks are virtually identical to those reported in the study by Derrfuss et al. (2005; but with slightly less agreement in the right hemisphere), where the authors also employed the ALE technique in one of its earlier implementations. Macro-anatomically, according to the Mindboggle 101 atlas (Klein et al. 2012; Manera et al. 2020), the LH FEF and RH FEF peaks lie within the caudal middle frontal gyrus (in BA 6; see Table 3) and not in the precentral gyrus, as previously assumed based on non-human primate evidence (Bruce et al 1985; Schall et al. 2020, and Tehovnik et al. 2000, for reviews). These results are consistent with the few pieces of evidence available on the delineation of this region based on cytoarchitecture in *post-mortem* studies (Rosano et al. 2003; Schmitt et al. 2005). While the left IFJ ALE peak was found in the precentral gyrus (in BA 6; see Table 3), interestingly the right IFJ ALE peak was instead localized within the pars opercularis (in BA 9). The distinctive architecture of the IFJ remains elusive, but these peaks agree with the general idea that this area is localized in several Brodmann areas (BA6, 8, 9, 44 and 45), and may correspond to a specific cyto- and chemo-architecture found dorsal to BA44 (Amunts and Von Cramon 2006; for a recent fine-grained analysis of its architecture see Ruland et al. 2022).

**Table 3.** Comparison of the ALE peaks with previous meta-analyses and brain atlases

| Cluster | Hemi | ALE peaks |  |  |  |  |  | Previous meta-analyses (Talairach space) |  |  |  |  |  | Brain atlases |  | LOEO results |
| --- | --- | --- | --- | --- | --- | --- | --- | --- | --- | --- | --- | --- | --- | --- | --- | --- |
|  |  | MNI152 coordinates |  |  | Talairach coordinates |  |  | Paus (1996) |  |  | Derrfuss et al. (2005) |  |  | Mindboggle 101 | MMP1 |  |
|  |  | x | y | z | x | y | z | x | y | z | x | y | z | Label | Label |  |
| FEF | L | -28 | -6 | 54 | -27.73 | -9.61 | 51.32 | -32 | -2 | 46 | // | // | // | Caudal Middle Frontal Gyrus | i6-8 | 26/35 |
| FEF | R | 30 | -6 | 50 | 27.17 | -9.86 | 48.03 | 31 | -2 | 47 | // | // | // | Caudal Middle Frontal Gyrus | 6a | 23/35 |
| IFJ | L | -42 | 6 | 30 | -40.87 | 3.27 | 30.37 | // | // | // | -40 | 4 | 30/32 | Precentral Gyrus | 6r | 31/32 |
| IFJ | R | 46 | 12 | 28 | 42.43 | 8.35 | 29.4 | // | // | // | 44 | 10 | 34 | Pars Opercularis | 6r | 24/32 |

Finally, in our opinion, the most interesting results of these comparisons were those obtained from the projection of our main FEF and IFJ clusters on the FSaverage surface using the method from Wu et al. (2017) where we could carefully examine their spatial relationship with the MMP1 (Glasser et al. 2016; see Figure 4). The FEF clusters covered almost the entire middle and anterior part of the FEF (as defined by the corresponding MMP1 label) but also large parts of the middle and posterior 6a region. Moreover, the left and right hemisphere ALE peaks were found within area i6-8 and area 6a, anteriorly and dorsally relative to the FEF, respectively. The IFJ clusters instead spanned multiple MMP1 labels, including areas PEF, 6r, IFJp and IFJa. While in the left hemisphere, the majority of the vertices of the cluster were localized in the middle and posterior aspect of the IFJp, in the right hemisphere most of the vertices were localized in the ventral 6r region. Crucially, in both hemispheres, however, the ALE peaks were localized in this region (i.e., 6r), ventrally relative to the IFJp.

**Fig. 4.**
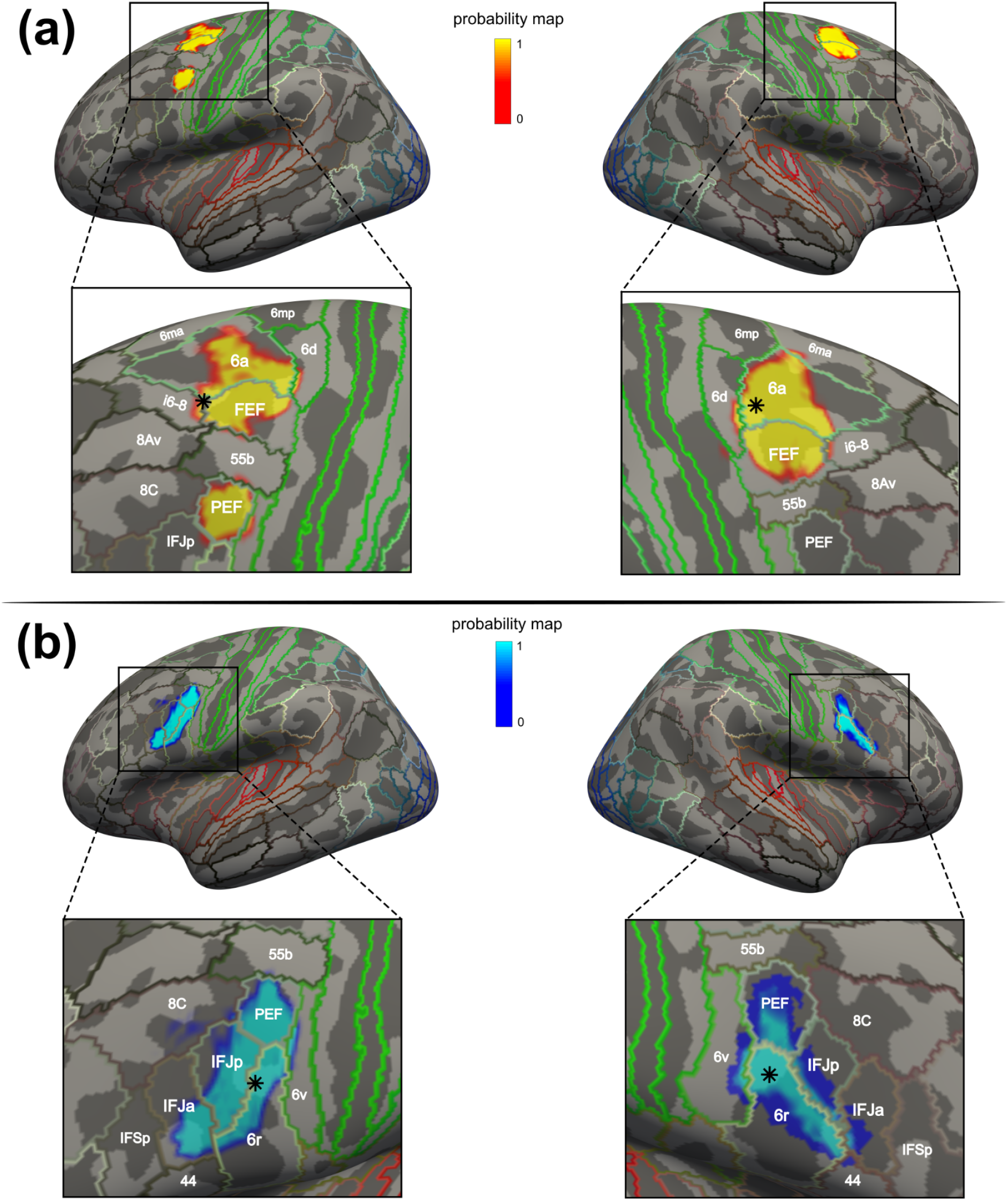
Projection on FSaverage of the FEF and IFJ main clusters and comparison with the MMP1 taxonomy. **A** Vertices corresponding to the FEF clusters. Both clusters covered the middle and rostral parts of the FEF label as defined according to the MMP1 atlas, but they also covered large parts of area 6a. In the left hemisphere, vertices were also localized in the iFEF, which matches almost exactly the boundaries of area PEF from the atlas. The LH FEF peak was localized within area i6-8, just anterior to the FEF, and the RH FEF peak was localized within area 6a, dorsal to FEF. Despite this difference, both peaks were localized near the junction of the sPCS and the SFS, in the anterior bank and the posterior banks of the sPCS, respectively. **B** Vertices corresponding to the IFJ clusters. They showed a similar elongated shape that approximately followed the posterior iPCS and encroached onto the IFS, and they spanned multiple MMP1 areas. Importantly, we found that in both hemispheres the IFJ peaks were localized near the junction of the iPCS and the SFS within area 6r, posteriorly to the IFJp

### Meta-analytic connectivity modeling results

The MACM analysis of the FEF and IFJ revealed a broad set of regions that coactivated with these seeds in the BrainMap database (see Figure 5) encompassing the frontal, parietal and temporal cortices. The LH FEF seed coactivated with other six clusters (see Figure 5, panel A), and the RH FEF coactivated with other eight clusters (Figure 5, panel B). Interestingly, while these FEF coactivations included as expected medial oculomotor regions (the SCEF) and the SPL/IPL, in both analyses we found coactivated clusters in the bilateral ventral PFC, which included parts of the iFEF and the IFJ based on their localization relative to the iPCS and the IFS. The LH IFJ coactivated with a broad set of other nine clusters (Figure 5, panel C), and the RH IFJ coactivated with only five other clusters (Figure 5, panel D). The coactivations of these bilateral seeds spread onto the IFS and ventrally in the insular cortex and claustrum. Again, these coactivations included clusters in the SCEF, the SPL/IPL and the angular gyrus. In contrast to the FEF coactivations, where the bilateral IFJ was always coactivated, we did not find FEF coactivations in the IFJ MACM results, except for the ipsilateral FEF in the LH IFJ MACM analysis. Another crucial difference was that in this analysis, we found a large cluster in the left temporal lobe that included the fusiform gyrus and the inferior occipital cortex. We confirmed these patterns by performing an ALE contrast analysis between the FEF and IFJ MACM results in each hemisphere (see Supplementary Information p. 16). To summarize, while the FEF MACM analysis showed that this region is consistently coactivated with the ventrolateral PFC and regions in the posterior parietal cortex across paradigms, the IFJ had more widespread coactivation patterns (particularly in the LH), being more strongly connected with the rest of the PFC and with the insular cortex and claustrum, and possessing a differential connectivity pattern with regions of the inferior temporal cortex. In contrast, we found differential coactivations of the LH and RH FEF with the ipsilateral precuneus/SPL and lateral intraparietal areas. In addition to revealing the task-based functional connectivity fingerprints of these regions (i.e., the coactivation patterns, as reported above), our functional decoding approach in BrainMap also allowed us to uncover the behavioral domains significantly associated with each of them (Figure 5, right side of each panel; see Supplementary Information p. 16 for a summary of these results).

**Fig. 5.**
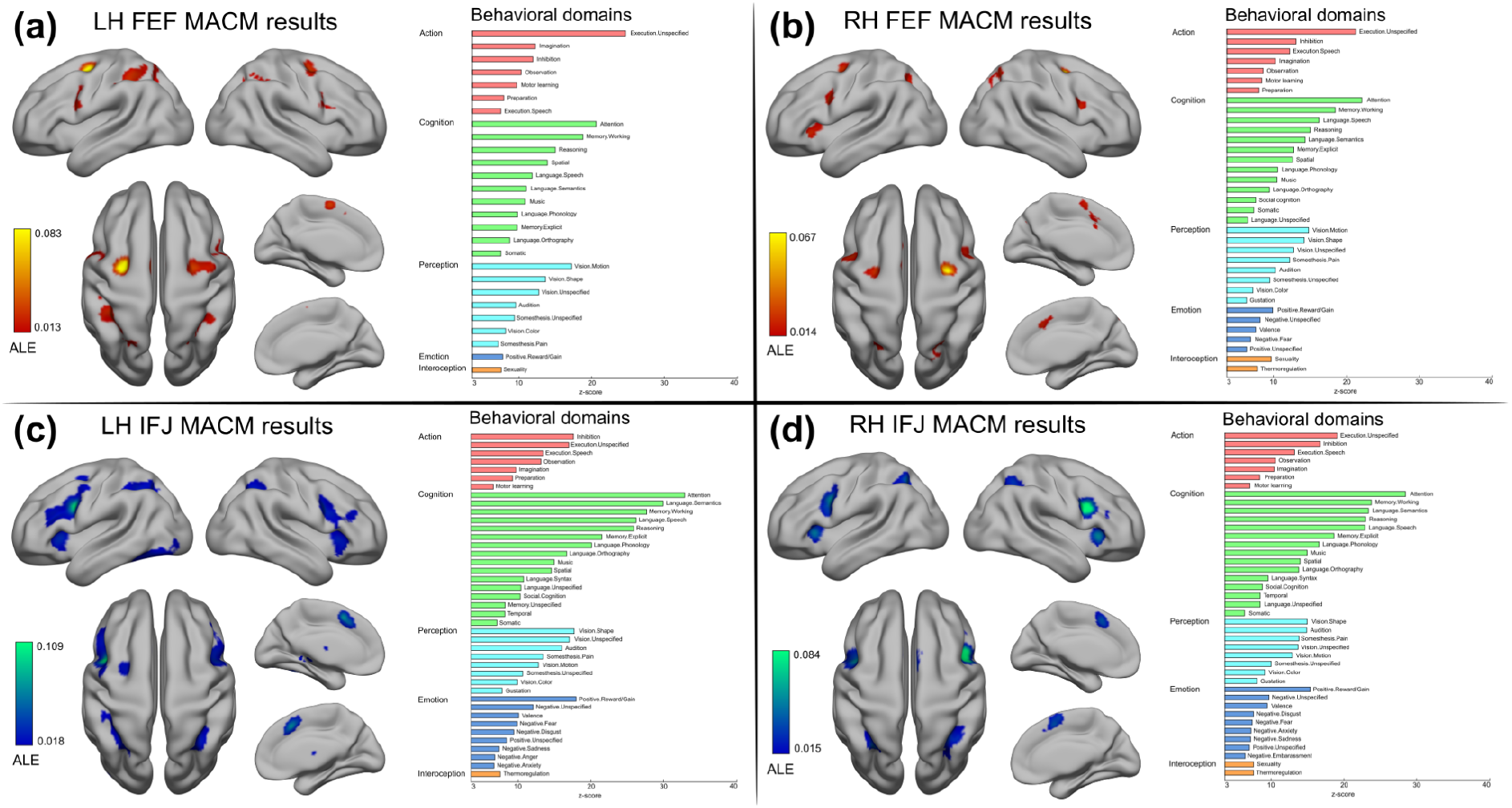
Meta-analytic connectivity modeling results. **A** and **B** depict the coactivation profiles of the FEF, and **C** and **D** the coactivation profiles of the IFJ. On the right side of each panel: decoding results of the significant associations with behavioral domains (p < 0.05, Bonferroni corrected)

## Discussion

The PFC is essential to several aspects of flexible goal-driven behavior that are mediated by specialized brain regions (Fuster 2000; Miller and Cohen 2001). The FEF and the IFJ have been previously implicated in covert spatial attention and oculomotor control on the one hand (Corbetta et al. 1998; Vernet et al. 2014), and working memory and cognitive control on the other (Bedini and Baldauf 2021; Brass et al. 2005). Their localization has been traditionally associated with the major sulci of the posterior-lateral PFC, namely the SFS and the sPCS, and the iPCS and the IFS, respectively (Brass et al. 2005; Paus 1996). Due to the large body of empirical work that has accumulated over the past years on these regions (Bedini and Baldauf 2021) and the parallel development of more robust meta-analytic techniques for neuroimaging data (Fox et al. 2014), we felt the need to reassess previous results in light of the current evidence, with a specific focus on overcoming discrepancies in the definition and localization of these regions using fMRI in humans. In particular, in this study, we sought to accurately estimate the precise localization of these regions in standard space by performing a coordinate-based meta-analysis using the ALE technique (Eickhoff et al. 2012). To model the spatial convergence of activations within the FEF, we analyzed data from 35 fMRI studies (35 experiments) that investigated the planning and execution of prosaccades and prosaccades and antisaccades contrasted against a fixation baseline in over 400 subjects. To model the spatial convergence within the IFJ, we analyzed data from 30 fMRI studies (32 experiments) that investigated visual attention, working memory, and cognitive control, in over 500 subjects. In contrast to previous ALE meta-analyses (Derrfuss et al. 2005: Grosbras et al. 2005: Jamadar et al. 2013) that relied on the false discovery rate, our study implemented a much more stringent method for multiple comparison correction (i.e., voxel-level FWE). We also included only 3T and 4T fMRI studies to make our inference spatially more reliable. Moreover, testing for significance in each voxel individually enabled us to carry out a fine-grained assessment of activations across experiments. Crucially, we found that by modeling activity across studies (thus partially overcoming inter-individual variability), sulcal landmarks are indeed consistently associated with both regions, as indicated by the location of the ALE peak values (see Figure 1 and Table 3). Our results thus suggest a robust association of structure and function in these higher regions (Frost and Goebel 2012; Miller et al. 2021; Van Essen 2007; Wang et al. 2015), similar to what previous studies have shown in early to mid-level visual regions (Benson et al. 2012; Hinds et al. 2008, 2009). We suggest that this association should be examined by future fMRI studies more systematically at the individual-subject level (Amiez and Petrides 2018; Derrfuss et al. 2009, 2012). Given the limitations of previous meta-analyses (Derrfuss et al. 2005; Grosbras et al. 2005: Jamadar et al. 2013; Paus 1996), we recommend using the coordinates reported in the present study to define FEF and IFJ based on the ALE technique.

The FEF is arguably one of the most important regions of a network involved in the planning and execution of saccadic eye movements and has been extensively studied in primates (Schall and Thompson 1999; Tehovnik et al. 2000). In humans, this network usually comprises a set of regions in the lateral and medial frontal cortex, posterior parietal cortex, and subcortical nuclei (Corbetta et al. 1998; Grosbras et al. 2005), and has been mainly investigated with fMRI over the past 25 years, enabling to characterize their respective functions (McDowell et al. 2008). Following the crucial foundation set by primate neurophysiology (Bruce et al. 1985; Buschman and Miller 2009; Moore and Fallah 2001), the human FEF has often been seen as not only implicated in the production of visually-guided and voluntary saccades and other oculomotor behaviors (Schall 2015), but also covert shifts of spatial attention, spatial working memory, and also more complex executive functions (Corbetta et al. 1998; Fiebelkorn and Kastner 2020; Vernet et al. 2014). Converging lines of research suggest that the FEF works as a spatial priority map (Fiebelkorn and Kastner 2020; Jerde et al. 2012; Sprague and Serences 2013). The localization of the human FEF is however highly debated and affected by strong spatial variability (see Bedini and Baldauf 2021, and Vernet et al. 2014 for a discussion), possibly due to inter-individual differences that are obscured when only reporting group-level results. Previous ALE meta-analyses provided evidence of consistent activations within FEF across PET and fMRI experiments investigating prosaccades and antisaccades (Cieslik et al. 2016; Grosbras et al. 2005; Jamadar et al. 2013). However, given the coarser spatial resolution of PET and low field strength MRI scanners (i.e., 1.5T) and acquisition methods that were included in these previous meta-analyses, these studies were only partially able to accurately infer spatial convergence across experiments, as well as dissociations across paradigms and contrasts. In addition, some of these studies relied on earlier implementations of the ALE technique, which allowed for within-study effects (see Turkeltaub et al. 2012), and critically, more liberal statistical thresholds and multiple comparisons correction solutions (such as false discovery rate, which is no longer recommended, as it is not considered appropriate for spatial inference; Eickhoff et al. 2016). Finally, another aspect that is difficult to evaluate retrospectively was the reporting of an implementation error in an earlier ALE version (Eickhoff et al. 2017), which affected the meta-analyses previous to that report. In this study, we attempted to overcome some of these limitations by applying a conservative thresholding method and by including higher field fMRI studies (i.e., 3T or 4T) to accurately infer the localization of the human FEF in standard space. Our ALE results, obtained by analyzing prosaccades > fixation, antisaccades > fixation, and prosaccades and antisaccades > fixation contrasts across 35 fMRI experiments (see Table S1), show that the highest spatial convergence based on the ALE value was found within two bilateral clusters in the dorsolateral PFC, localized in the anterior bank of the sPCS, near its junction with the SFS (see Figure 1; Table 2). The comparison with one of the most comprehensive brain parcellations available to date, namely the MMP1 (Glasser et al. 2016), revealed some interesting topographic differences. While many of the projected voxels of our FEF clusters were localized within the middle and anterior part of the MMP1 FEF region, a considerable portion was also spreading dorsally in the middle and posterior 6a region. Furthermore, the ALE peaks were localized within area i6-8 in the left hemisphere, and area 6a in the right hemisphere according to the MMP1. In the left hemisphere, a ventral cluster corresponding to the inferior or lateral FEF (Derrfuss et al. 2012; Kastner et al. 2007; Luna et al. 1998) overlapped almost entirely with the PEF. These results suggest that there may be important differences in the way the FEF is defined across methods. The MMP1 was created by a careful manual delineation combining structural MRI (cortical thickness and myelin ratio), resting-state fMRI connectivity, and retinotopic mapping techniques (Glasser et al. 2016). Additionally, fMRI contrasts from nine tasks were also employed to infer areal boundaries (Barch et al. 2013), which were chosen to optimally balance breadth vs depth and scan time (Elam et al. 2021). Although we regard the MMP1 as a step change in our understanding of brain organization, and in particular of the fine-grained organization and structure of the PFC both in humans and in non-human primates (Donahue et al. 2018), we would like to suggest that more information gathered from task-based fMRI will be needed to better understand the functional subdivision of the plPFC. In particular, following the taxonomy proposed by the MMP1, major efforts should be made to isolate FEF activity from posterior activity in the premotor cortex on the one hand (area 6d), and from a newly discovered language selective region that borders the FEF ventrally on the other, namely area 55b (for a discussion on inter-individual variability in the spatial organization of these regions see Glasser et al. 2016). Ultimately, future developments of a functional localization method will facilitate the convergence of atlas-based and fMRI information to allow the delineation of anatomical clusters of activation within FEF with adequate functional specificity. Our ALE results partly suggest that this is needed as the current gold standard for localizing the FEF may lead to the inclusion of voxels from heterogeneous brain regions as defined according to the MMP1 atlas, possibly also due to the greater inter-subject variability that characterizes the posterior-lateral PFC (Bedini and Baldauf 2021; Glasser et al. 2016).

In this direction, a very compelling set of results were reported by Mackey et al. (2017), who identified two retinotopic maps in the sPCS (sPCS1 and SPC2) and a third one in the iPCS. By examining the correspondences between their results and the MMP1 (see Figure 8 from their study), they found that the sPCS2 corresponded to the FEF, while the sPCS1 corresponded to areas 6a and 6d. Interestingly, they also reported that in all subjects and both hemispheres, the foveal representation was localized in the fundus of the sPCS, at its intersection with the SFS. This description closely matches the localization of our ALE peaks, which raises the question of whether the fMRI contrasts we included in our meta-analysis could be targeting specific neural populations within the FEF. It is well established that in the macaque, a population of neurons shows increased firing rates when the animal is fixating and is inhibited when executing saccades (hence termed ‘fixation’ neurons; Bruce and Golberg 1985; Hanes et al. 1998; Lowe and Schall 2018). Are these populations of neurons also present in humans, and how are they distributed within the FEF? Are these neurons associated with the significant increase in the BOLD signal when comparing saccades to peripheral positions against a fixation baseline, and what is the role of saccadic amplitude in isolating peaks of activity within the FEF (see Grosbras 2016)? An additional aspect that may be worth investigating is whether the activations found in one or more of these clusters (for example, the iFEF) are dependent on some artifacts present in the experimental design or analysis. In the 35 experiments we analyzed, 10 didn’t record eye movements in the scanner (see Table S1), making it at least likely that some of these clusters were driven by mixed signals and in the worst case, by spurious neural activity that was not related to saccadic behavior. For example, it is well documented that eye blinks can contaminate BOLD signal (Bristow et al. 2005; Hupé et al. 2012), and this fact was invoked to explain discrepancies in the oculomotor organization in primates (Tehovnik et al. 2000) and as a signal driving iFEF responses (Amiez and Petrides 2009; Kato and Miyauchi 2003). In conclusion, we strongly agree with the general caveat that the way the FEF is defined is ultimately constrained by the technique employed (Schall et al. 2020; Vernet et al. 2014), and in particular its spatial resolution. The localization and the extent of the FEF cluster should be inferred based on the convergence of multiple criteria (primarily architectonic, sulcal, functional, connectional, and also comparative). With the present ALE meta-analysis, we aimed at providing updated quantitative evidence on the localization of this region in standard space using standard functional localizers across over 400 subjects. However, additional efforts will be needed to understand the organization of the human eye fields in the PFC and their relationship with sulcal neuroanatomy at the individual-subject level (Amiez et al. 2006; Amiez and Petrides 2018; Frost and Goebel 2012). Careful mapping of the localization of the human FEF is also essential to bridge research in humans and non-human primates and will be crucial for testing hypotheses about homologies in the organization of the prefrontal cortex across species, for example, based on connectivity information (Mars et al. 2021; Neggers et al. 2015; Sallet et al. 2013).

The study of the role of the ventrolateral PFC in various cognitive functions such as non-spatial attention, working memory and cognitive control led to the definition of the IFJ as a separate brain region involved in critical aspects of all these functions (Brass et al. 2005; Derrfuss et al. 2004, 2005). This region seems to be tightly coupled with specific sulcal landmarks across individuals, namely the junction of the iPCS and the IFS (Derrfuss et al. 2009) and belongs to the frontoparietal network (Cole and Schneider 2007; Cole et al. 2013; Yeo et al. 2011). Recent fMRI evidence from a large sample moreover showed that the IFJp is one of the core multiple demand regions of the brain (Assem et al. 2020). In this study, we therefore pooled together results from the various tasks that have been used to localize this region (see Table S2) ranging from attentional (i.e., RSVP/oddball; Asplund et al. 2010; and endogenous cueing paradigms; Baldauf and Desimone 2014; Zhang et al. 2018), working memory (primarily n-back paradigms; Zanto et al. 2010), and cognitive control paradigms (i.e., task-switching and Stroop tasks; Brass and Von Cramon 2002; Derrfuss et al. 2012). Following a previous meta-analysis (Derrfuss et al. 2005), we reasoned that the spatial convergence across these paradigms (rather than a single task) would allow us to infer the accurate localization of the IFJ in standard space. Consistent with our hypothesis, we found two prominent clusters of activation in the ventral PFC. Based on the respective ALE peak values, the highest convergence was found in the posterior bank of the iPCS, approximately at the height of its ventral junction with the IFS (see Figure 3; Table 2). The comparison of these results with the MMP1 (Glasser et al. 2016) through their projection to the FSaverage surface revealed additional interesting topographic differences. Both IFJ clusters spanned several regions of the MMP1, including areas PEF, 6r, IFJp, and IFJa (Figure 6, panel B), with the majority of the voxels being localized around the borders of IFJp, 6r and the PEF. Moreover, both ALE peaks were localized in area 6r, ventral to the IFJp. These results suggest that the paradigms currently employed to localize the IFJ, and some of those which can be considered promising candidates for this purpose, will also tend to activate voxels from several other brain regions as defined according to this atlas. This problem is further exacerbated by the fact that the IFJp itself and the neighboring regions are also part of the multiple demand system, so many experimental manipulations will tend to coactivate some of them at the same time, thus concealing its boundaries. We examined potential differences in localization between paradigms in a control analysis (see Figure S6 in the Supplementary Information). Taken together, our results showed that oddball/attention paradigms tend to activate voxels that are localized in the posterior aspect of the IFS, whereas both working memory and cognitive control paradigms tend to activate more posterior voxels in the banks of the iPCS. These results seem to suggest that these paradigms consistently recruit different IFJ subregions and additional regions, although their overlap is precisely localized within the IFJ.

An interesting approach that recognizes the integrative nature of the IFJ draws from the combination of different paradigms and uses the conjunction of the respective activations to localize this region, which may overcome some of the previous limitations. Stiers and Goulas (2018) analyzed the voxel responses across three different tasks (Eriksen flanker task, backmatching or n-back task, and a response switching task) to define the prefrontal nodes of the multiple demand system in 12 subjects. A manipulation of task difficulty in each of the previous tasks was used to identify voxels that were modulated by increasing cognitive demands, which were used to define ROIs in each subject in a conjunction analysis across tasks for further analyzing their relative task preference and functional connectivity patterns. Their conjunction analysis revealed local maxima of activity within the IFJ, where voxels with different task preferences exhibited distinct functional connectivity patterns with the rest of the brain (Stiers and Goulas 2018). Based on these results, it may be argued that no single task alone would adequately capture the selectivity patterns of neural populations within the IFJ; rather, manipulations of task difficulty combined with the administration of different tasks could provide an unbiased way of localizing this region in individual participants. The results of the present study aimed at providing quantitative evidence of the convergence of activations within the IFJ across non-spatial attentional, working memory, and cognitive control tasks across over 500 subjects, which we hope will help guide future research aimed at understanding the relationship between the function and localization of the IFJ, and its relationship with sulcal neuroanatomy at the individual subject level (Derrfuss et al. 2009). Future meta-analyses should better clarify how activations from different tasks that tap cognitive control functions map onto the ventro-lateral PFC and assess whether they may recruit distinct subregions near the IFJ (see for example Kim et al. 2012, and Nee et al. 2013), which was however outside the scope of our present work.

A secondary goal of the present study was to uncover the task-based functional connectivity fingerprint (Passingham et al. 2002) of the FEF and the IFJ in a data-driven fashion. We retrieved in the BrainMap database all the studies that reported activations within a cuboid seed centered around the FEF and IFJ standard coordinates found in our ALE main localizer analyses and we employed the MACM technique (Langner and Camilleri 2021) to uncover their coactivation profiles. Importantly, while previous studies already performed MACM analyses on the FEF (Cieslik et al. 2016) and the IFJ (Muhle-Karbe et al. 2016; Sundermann and Pfleiderer 2012), our study is to our knowledge the first that used this technique on the results of an ALE analysis specifically aimed at localizing these regions (and not a manual or atlas-based delineation) using a conservative seed extent (6 mm). Our MACM analysis allowed adequately powered inference in each seed region (n > 17 experiments; Eickhoff et al. 2016) and revealed broad networks of coactivations that encompassed the frontal, parietal and temporal cortices (see Figure 5). The most remarkable differences between FEF and IFJ coactivation patterns were that on the one hand, the LH and RH FEF coactivated with the bilateral ventrolateral PFC (iFEF and IFJ), whereas only the LH IFJ coactivated with the left FEF in the experiments retrieved. On the other hand, the LH IFJ had stronger and more widespread coactivations in PFC and the insular cortex and was also coactivated with the inferotemporal cortex. These coactivation patterns may be essential for the IFJ to perform its role in feature- and object-based attention tasks (Baldauf and Desimone 2014; Gong and Liu 2020; Meyyappan et al. 2021) and could be in turn supported by its underlying anatomical connectivity patterns (Baldauf and Desimone 2014; Bedini et al. 2021). These results are consistent with the idea of a dorso-ventral segregation of fronto-parietal coactivations forming a spatial/motor and a non-spatial/motor network, and that is in turn associated with the first and third branch of the superior longitudinal fasciculus, respectively (Parlatini et al. 2017). In addition, our functional decoding results suggest that these systematic coactivation patterns allow these regions to support multiple yet specialized roles in flexible goal-driven behavior (Assem et al. 2020; Fiebelkorn and Kastner 2020; Ngo et al. 2019).

## Conclusion

Our study provides the accurate localization of two regions of the posterior lateral PFC, namely the FEF and the IFJ. These regions are tightly linked to sulcal landmarks as measured using fMRI across over 400 hundred subjects, with the FEF being localized at the junction of the sPCS and the SFS, and the IFJ at the junction of the iPCS and the IFS. Functionally, they appear to be organized according to a dorso-ventral gradient, going from areas responsible for sensorimotor transformations and action execution (FEF, iFEF), to areas that are involved in maintaining and updating behavioral goals according to internal representations (IFJ; Nee et al. 2013; O’Reilly 2010). Taken together, our findings aim at proposing a consensus standard localization of these regions in standard space, and meta-analytic groundwork to investigate the relationship between functional specialization and connectivity in large publicly available neuroimaging datasets (e.g., Markiewicz et al. 2021; Van Essen et al. 2013), as well as to guide future non-invasive brain stimulation studies.

## Author contributions

Conceptualization: **MB**, **DB**; Methodology: **MB**, **EO**, **PA**, **DB**; Software: **MB**; Validation: **MB**; Formal analysis; **MB**; Investigation: **MB**; Resources: **MB**, **DB**; Data curation: **MB**; Writing - original draft: **MB**; Writing - review and editing: **MB**, **EO**, **PA**, **DB**; Visualization: **MB**; Supervision: **EO**, **PA**, **DB**; Project administration: **MB**; Funding acquisition: **MB**.

## Data availability

The authors are committed to making the datasets generated during the current study available in a Neurovault repository upon article acceptance.

## Acknowledgments

We would like to thank Luca Turella and Daniel Adams for their comments on the preliminary results of this study. We would also like to thank Gabriele Amorosino and Francesca Saviola for their technical advice, and Vittorio Iacovella and Giorgio Marinato for their advice on resource/data sharing.

