## Supplementary_Information for "Accurate localization and coactivation profiles of the Frontal Eye Field and Inferior Frontal Junction: an ALE and MACM fMRI meta-analysis"

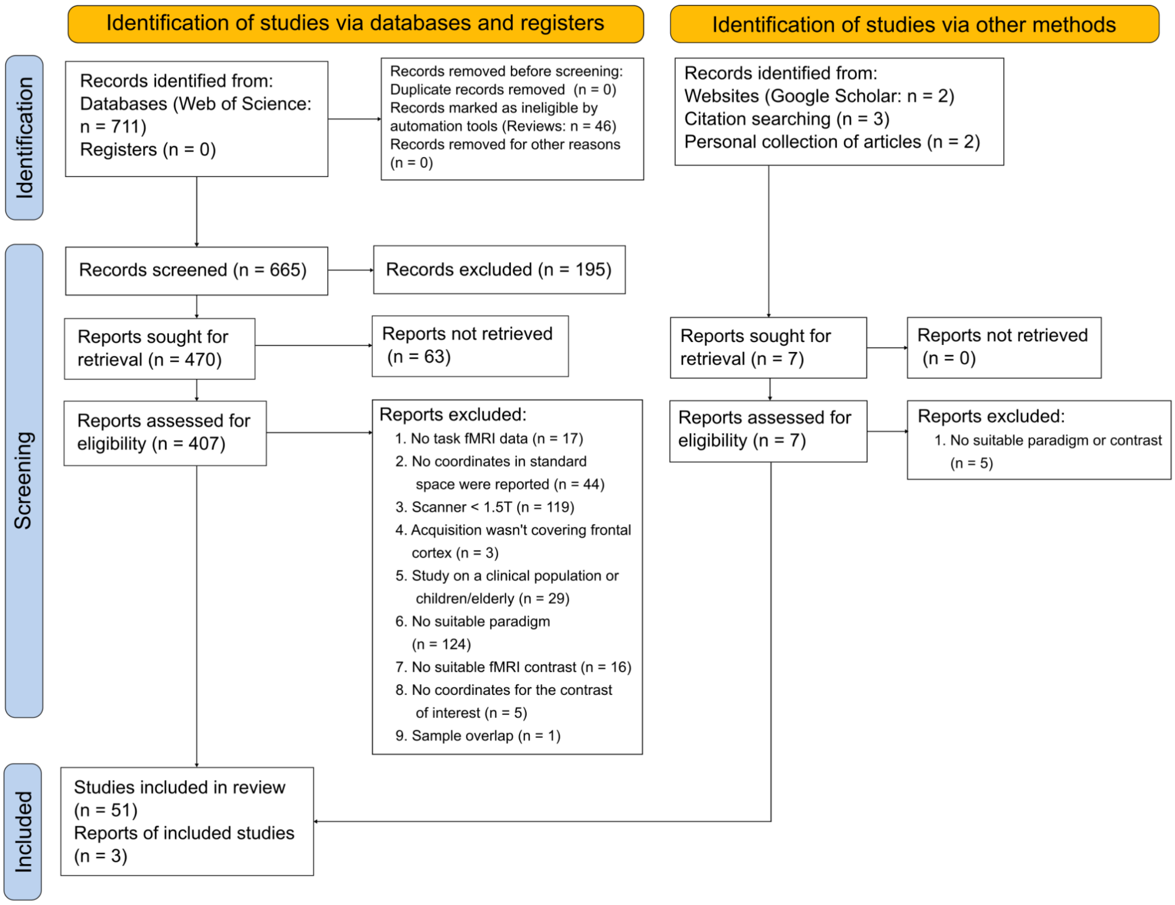


**Fig. S1** PRISMA flow chart FEF sample

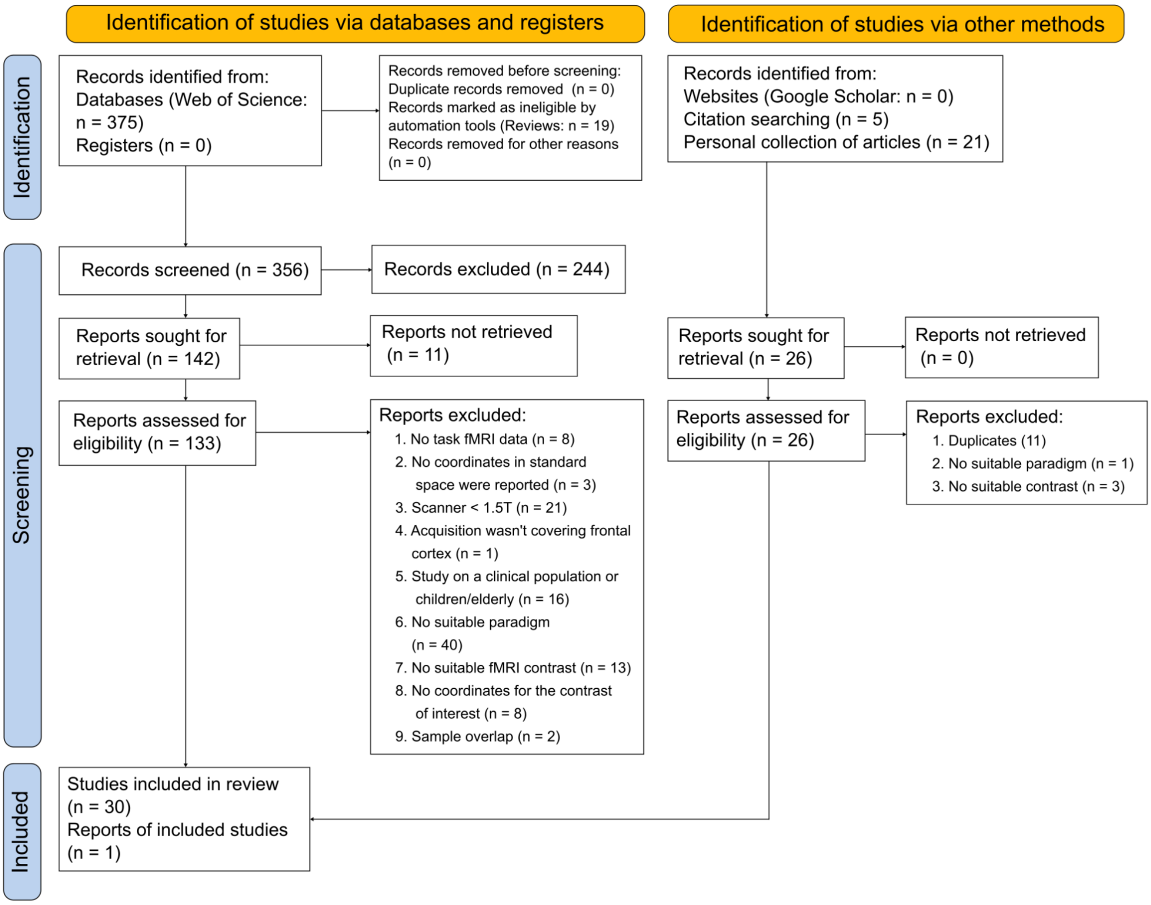


**Fig S2** PRISMA flow chart IFJ sample

**Supplementary Table 1** List of the studies included in the FEF localizer sample

| Study | N | Age | Paradigm | Contrast | Scanner | Design | Eye tracker | Independent fMRI localizer | Field of View (FOV) | Results reported for | N° of foci | Multiple FEF foci | Software | Space |
|---|---|---|---|---|---|---|---|---|---|---|---|---|---|---|
| Alkan et al. (2011) | 8 | 26 ± 4 | Oculomotor task | Prosaccades > Fixation | 3T | Blocked | Y | N | 220 mm | Whole-brain | 19 | N | AFNI | Talairach |
| Amiez and Petrides (2018) | 13 | 22.6 ± 2.8 | Oculomotor task | Prosaccades > Fixation | 3T | Blocked | N | N | NA | FEF foci | 2 | N | SPM12b | MNI |
| Atmaca et al. (2013) | 11 | 26.9; 22-33 | Oculomotor task | Prosaccades > Fixation | 3T | Blocked | N | Y | 192 mm | Whole-brain | 24 | Y | SPM8 | MNI |
| Bär et al. (2016) | 14 | 22-56 | Oculomotor task | Prosaccades > Fixation | 3T | Blocked | N | N | 240 mm | Whole-brain | 12 | N | BrainVoyager QX 1.10 | Talairach |
| Bär et al. (2016) | 14 | 22-56 | Oculomotor task | Antisaccades > Fixation | 3T | Blocked | N | N | 240 mm | Whole-brain | 18 | Y | BrainVoyager QX 1.10 | Talairach |
| Berman et al. (1999) | 11 | 25.6 ± 7.1; 18-43 | Oculomotor task | Prosaccades > Fixation | 3T | Blocked | N | N | 40 x 20 mm | ROIs | 14 | Y | AFNI | Talairach |
| Braga et al. (2016) | 20 | 26.2; 21-36 | Oculomotor task | Prosaccades > Fixation | 3T | Blocked | Y | Y | 220 mm | Whole-brain | 7 | N | FSL | MNI |
| Brown et al. (2006) | 10 | 26; 22-33 | Oculomotor task | Prosaccades > Fixation | 4T | *Event-related* | Y | N | 220 mm | Whole-brain | 17 | N | BrainVoyager 2000 | Talairach |
| Brown et al. (2006) | 10 | 26; 22-33 | Oculomotor task | Antisaccades > Fixation | 4T | *Event-related* | Y | N | 220 mm | Whole-brain | 21 | N | BrainVoyager 2000 | Talairach |
| Connolly et al. (2000) | 7 | 24.8 ± 3.2 | Oculomotor task | Prosaccades > Fixation | 4T | Blocked | N | N | 192 mm | Whole-brain | 7 | Y | Stimulate | Talairach |
| Connolly et al. (2000) | 7 | 24.8 ± 3.2 | Oculomotor task | Antisaccades > Fixation | 4T | Blocked | N | N | 192 mm | Whole-brain | 13 | Y | Stimulate | Talairach |
| Connolly et al. (2002) | 8 | NA | Oculomotor task | Prosaccades > Fixation | 4T | Blocked | N | Y | 192 mm | ROIs | 4 | N | Stimulate | Talairach |
| Connolly et al. (2005) | 5 | NA | Oculomotor task | Prosaccades > Fixation | 4T | Blocked | Y | Y | 192 mm | ROIs | 5 | N | Stimulate / Brain Voyager 4.9 | Talairach |
| Connolly et al. (2007) | 8 | NA | Oculomotor task | Prosaccades > Fixation | 4T | Blocked | N | Y | 192 mm | ROIs | 2 | Y | Stimulate / BrainVoyager 4.9 / QX | Talairach |
| Christophel et al. (2018) | 22 | 24.4 ± 0.83 | Oculomotor task | Prosaccades > Fixation | 3T | *Event-related* | N | Y | 192 mm | FEF foci | 2 | N | SPM8 | MNI |
| Curtis & Connolly (2008) | 12 | 21-35 | Oculomotor task | Pro- & Antisaccades > Fixation | 3T | *Event-related* | Y | N | 192 mm | ROIs | 32 | Y | Caret | MNI |
| DeSouza et al. (2003) | 10 | 26.6 ± 1.0 | Oculomotor task | Pro- & Antisaccades > Fixation | 4T | Blocked | Y | Y | 192 mm | Whole-brain | 9 | N | BrainVoyager 2000 4.4 | Talairach |
| Duecker et al. (2013) | 20 | 19-28 | Oculomotor task | Prosaccades > Fixation | 3T | Blocked | N | Y | 192 mm | FEF foci | 2 | N | BrainVoyager QX 2.3 | Talairach |
| Fernandez-Ruiz et al. (2018) | 25 | 21.7 ± 1.9; 18-25 | Oculomotor task | Pro- & Antisaccades > Fixation | 3T | *Event-related* | Y | N | 211 mm | Whole-brain | 18 | N | Brain Voyager QX 2.8.4 | Talairach |
| Frannson et al. (2014) | 24 | 26.7 ± 4.9 | Oculomotor task | Prosaccades > Fixation | 3T | Blocked | Y | N | 288 mm | Whole-brain | 8 | N | SPM8 | MNI |
| Furlan et al. (2016) | 6 | NA | Oculomotor task | Pro- & Antisaccades > Fixation | 3T | *Event-related* | Y | N | 192 mm | ROIs | 6 | N | BrainVoyager QX 2.3 | Talairach |
| Guo et al. (2012) | 12 | 19-31 | Oculomotor task | Prosaccades > Fixation | 3T | Blocked | Y | Y | 192 mm | ROIs | 6 | N | SPM8 | Talairach |
| Gurel et al. (2018) | 16 | 23.2 | Oculomotor task | Prosaccades > Fixation | 3T | Blocked | Y | Y | 192 mm | FEF foci | 2 | N | BrainVoyager QX | Talairach |
| Heinen et al. (2006) | 5 | 20-37 | Oculomotor task | Prosaccades > Fixation | 3T | Blocked | Y | Y | NA | ROIs | 4 | N | VISTASOFT | Talairach |
| Hubl et al. (2008) | 7 | 31 ± 9 | Oculomotor task | Prosaccades > Fixation | 3T | Blocked | N | Y | 192 mm | Whole-brain | 9 | N | BrainVoyager QX | Talairach |
| Jamadar et al. (2015) | 23 | 25.8; 18-43 | Oculomotor task | Prosaccades > Fixation | 3T | *Event-related* | Y | Y | 192 mm | Whole-brain | 47 | N | SPM8 | MNI |
| Jamadar et al. (2015) | 23 | 25.8; 18-43 | Oculomotor task | Antisaccades > Fixation | 3T | *Event-related* | Y | Y | 192 mm | Whole-brain | 47 | N | SPM8 | MNI |
| Jarvstad & Gilchrist (2019) | 23 | NA | Oculomotor task | Prosaccades > Fixation | 3T | Blocked | Y | N | 192 mm | Whole-brain | 4 | N | FSL v.5.06-1 | MNI |
| Kastner et al. (2007) | 4 | 20-36 | Oculomotor task | Prosaccades > Fixation | 3T | Blocked | Y | Y | 256 mm | ROIs | 5 | N | AFNI | Talairach |
| Kurata et al. (2005) | 6 | 29-38 | Oculomotor task | Prosaccades > Fixation | 3T | Blocked | N | N | 220 mm | Whole-brain | 14 | N | Advanced Visual Systems & AFNI | Talairach |
| Levy et al. (2007) | 4 | 24-43 | Oculomotor task | Prosaccades > Fixation | 3T | Blocked | N | Y | 192 mm | ROIs | 5 | N | BrainVoyager QX | Talairach |
| Neggers et al. (2007) | 15 | NA | Oculomotor task | Pro- & Antisaccades > Fixation | 3T | Blocked | N | Y | 224 x 256 x 128 mm | ROIs | 4 | Y | SPM2 | MNI |
| Pierce et al. (2019) | 30 | 25.7 ± 4.1 | Oculomotor task | Prosaccades > Fixation | 3T | Blocked | Y | Y | NA | Whole-brain | 9 | Y | SPM12 | MNI |
| Schon et al. (2008) | 17 | 21.29 ± 3.72 | Oculomotor task | Prosaccades > Fixation | 3T | Blocked | N | Y | 200 mm | Whole-brain | 19 | N | SPM2 | MNI |
| Schwerdtfeger et al. (2013) | 14 | 29.6 ± 9.6 | Oculomotor task | Pro- & Antisaccades > Fixation | 3T | *Event-related* | Y | N | 211 x 211 mm | ROIs | 10 | Y | Brain Voyager QX 1.9 | Talairach |
| Tamber-Rosenau et al. (2018) | 8 | 28.5 ± 3.3 | Oculomotor task | Prosaccades > Fixation | 3T | Blocked | Y | Y | 240 x 240 mm | ROIs | 6 | N | BrainVoyager QX v.1.10.2–2.8 | Talairach |
| Tark & Curtis (2009) | 5 | 22-39 | Oculomotor task | Prosaccades > Fixation | 3T | Blocked | Y | Y | 192 mm | FEF foci | 2 | N | Caret | MNI |
| Tibber et al. (2010) | 16 | 21-40 | Oculomotor task | Prosaccades > Fixation | 3T | Blocked | Y | Y | 192 mm | ROIs | 9 | N | SPM5 | Talairach |
| Tu et al. (2006) | 10 | 27.9 ± 3.18 | Oculomotor task | Antisaccades > Fixation | 3T | Blocked | N | Y | 250 x 250 mm | Whole-brain | 21 | N | SPM2 | MNI |

**Supplementary Table 2** List of the studies included in the IFJ localizer sample

| Study | N | Age | Paradigm | Contrast | Scanner | Design | Eye tracker | Independent fMRI localizer | Field of View (FOV) | Results reported for | N° of foci | IFJ's activity lateralization | Software | Space |
|---|---|---|---|---|---|---|---|---|---|---|---|---|---|---|
| Armbruster et al. (2012) | 20 | 23.5; 20–32 | Task-switching paradigm | Task switch > Distractor trials | 3T | *Event-related* | N | N | 192 mm | Whole-brain | 15 | Bilateral | SPM8 | MNI |
| Armbruster et al. (2012) | 20 | 23.5; 20–32 | Task-switching paradigm | Distractor inhibition > Baseline | 3T | *Event-related* | N | N | 192 mm | Whole-brain | 9 | Bilateral | SPM8 | MNI |
| Armbruster et al. (2012) | 20 | 23.5; 20–32 | Task-switching paradigm | Task switch > Repetition trials | 3T | *Event-related* | N | N | 192 mm | Whole-brain | 13 | Left | SPM8 | MNI |
| Asplund et al. (2010) | 30 | NA | RSVP / Oddball paradigm | Surprise > Search trials | 3T | *Event-related* | N | N | 240 mm | Whole-brain | 17 | Bilateral | BrainVoyager 4.9.1, QX 1.7.9 | Talairach |
| Asplund et al. (2010) | 30 | NA | RSVP / Oddball paradigm | Search trials > Baseline | 3T | *Event-related* | N | N | 240 mm | Whole-brain | 8 | Bilateral | BrainVoyager 4.9.1, QX 1.7.9 | Talairach |
| Asplund et al. (2010) | 6 | NA | RSVP / Oddball paradigm | Search trials > Baseline | 3T | *Event-related* | N | N | 240 mm | ROIs | 8 | Bilateral | BrainVoyager QX 1.11.4 | Talairach |
| Asplund et al. (2010) | 6 | NA | RSVP / Oddball paradigm | Surprise > Search trials | 3T | *Event-related* | N | N | 240 mm | ROIs | 8 | Bilateral | BrainVoyager QX 1.11.4 | Talairach |
| Baldauf and Desimone (2014) | 12 | 23-37 | Object-based attention paradigm | Attend face/house blocks > Difficulty-mathced rare target detection | 3T | Blocked | N | Y | N/A | IFJ only | 2 | Bilateral | SPM8 | MNI |
| Bode and Haynes (2009) | 12 | 26.4; 24-30 | Task-switching paradigm | Cue presentation > Baseline | 3T | *Event-related* | N | N | N/A | Whole-brain | 7 | NA | SPM2 | MNI |
| Bode and Haynes (2009) | 12 | 26.4; 24-30 | Task-switching paradigm | Target presentation > Baseline | 3T | *Event-related* | N | N | N/A | Whole-brain | 12 | NA | SPM2 | MNI |
| Bode and Haynes (2009) | 12 | 26.4; 24-30 | Task-switching paradigm | Response > Baseline | 3T | *Event-related* | N | N | N/A | Whole-brain | 9 | NA | SPM2 | MNI |
| Bollinger et al. (2010) | 18 | 23.4 ± 3.06; 18-28 | Working memory paradigm | FC with FFA: Stimulus known > Stimulus unknown | 3T | *Event-related* | N | N | 230 mm | Whole-brain | 17 | Right | SPM5 | MNI |
| Bollinger et al. (2010) | 18 | 23.4 ± 3.06; 18-28 | Working memory paradigm | FC with FFA: Stimulus known > Passive view + stimulus unknown | 3T | *Event-related* | N | N | 230 mm | Whole-brain | 17 | Right | SPM5 | MNI |
| Brass and Von Cramon (2002) | 11 | 26.2 ± 3.06 | Task-switching paradigm | Cue-only trials > Null events | 3T | *Event-related* | N | N | 192 mm | Whole-brain | 22 | Bilateral | LIPSIA | Talairach |
| Brass and Von Cramon (2002) | 11 | 26.2 ± 3.06 | Task-switching paradigm | Cue-target > Cue-only trials | 3T | *Event-related* | N | N | 192 mm | Whole-brain | 21 | Left | LIPSIA | Talairach |
| Brass and Von Cramon (2002) | 11 | 26.2 ± 3.06 | Task-switching paradigm | Cue-target trials > Null events | 3T | *Event-related* | N | N | 192 mm | Whole-brain | 10 | Bilateral | LIPSIA | Talairach |
| Brass and Von Cramon (2002) | 11 | 26.2 ± 3.06 | Task-switching paradigm | Cue-target trials > No-cue-target trials | 3T | *Event-related* | N | N | 192 mm | Whole-brain | 9 | Bilateral | LIPSIA | Talairach |
| Brass and Von Cramon (2004) | 14 | 24.4 ± 1.9 | Task-switching paradigm | Meaning switch > Cue switch trials | 3T | *Event-related* | N | N | 192 mm | ROIs | 3 | Left | LIPSIA | Talairach |
| Brass and Von Cramon (2004) | 14 | 24.4 ± 1.9 | Task-switching paradigm | Cue switch > Cue repetition trials | 3T | *Event-related* | N | N | 192 mm | ROIs | 4 | NA | LIPSIA | Talairach |
| Brass and Von Cramon (2004) | 14 | 24.4 ± 1.9 | Task-switching paradigm | Switch > Repetition trials (single cue) | 3T | *Event-related* | N | N | 192 mm | ROIs | 3 | Left | LIPSIA | Talairach |
| Chen et al. (2013) | 23 | 21 ± 1.67 | Stroop paradigm | Stimulus incongruent > Congruent trials (early stage) | 3T | *Event-related* | N | N | 250 x 250 mm | Whole-brain | 10 | NA | SPM8 | Talairach |
| Chen et al. (2013) | 23 | 21 ± 1.67 | Stroop paradigm | Response incongruent > Stimulus incongruent (early stage) | 3T | *Event-related* | N | N | 250 x 250 mm | Whole-brain | 5 | NA | SPM8 | Talairach |
| Chen et al. (2013) | 23 | 21 ± 1.67 | Stroop paradigm | Response incongruent > Stimulus incongruent (late stage) | 3T | *Event-related* | N | N | 250 x 250 mm | Whole-brain | 9 | NA | SPM8 | Talairach |
| Cole and Schneider (2007) | 9 | 19-42 | Working memory paradigm | Target switching > Non-switching trials (target non-occluded) | 3T | Mixed blocked/event-related | N | N | 210 mm | Whole-brain | 12 | Bilateral | BrainVoyager QX | Talairach |
| Cole and Schneider (2007) | 9 | 19-42 | Working memory paradigm | Target switching > Non-switching trials (target occluded) | 3T | Mixed blocked/event-related | N | N | 210 mm | Whole-brain | 5 | NA | BrainVoyager QX | Talairach |
| Corradi-Dell'Acqua et al. (2015) | 18 | 28; 18-52 | Object-based attention paradigm | High saliency unattended > attended stimuli | 3T | *Event-related* | N | N | N/A | Whole-brain | 5 | Bilateral | SPM8 | MNI |
| Derrfuss and Von Cramon (2004) | 19 | 20-36 | Task-switching paradigm | Switch > Null event trials | 3T | *Event-related* | N | N | 192 mm | IFJ only | 2 | Bilateral | LIPSIA | Talairach |
| Derrfuss and Von Cramon (2004) | 19 | 20-36 | Stroop paradigm | Incongruent > Neutral trials | 3T | *Event-related* | N | N | 192 mm | IFJ only | 2 | Bilateral | LIPSIA | Talairach |
| Derrfuss and Von Cramon (2004) | 19 | 20-36 | Working memory paradigm | 2-back > 0-back blocks | 3T | Blocked | N | N | 192 mm | IFJ only | 2 | Bilateral | LIPSIA | Talairach |
| Derrfuss et al. (2012) | 12 | 25.3 ± 2.4; 22-31 | Stroop paradigm | Incongruent > Congruent trials | 3T | *Event-related* | N | N | 220 x 220 mm | IFJ only | 1 | Left | FSL | MNI |
| Han and Marois (2013) | 14 | 20-32 | RSVP paradigm | Discontinuous > Continuous & continuous-hard conditions | 3T | *Event-related* | N | Y | 240 mm | Whole-brain | 14 | Bilateral | Brain Voyager QX 1.10 | Talairach |
| Han and Marois (2014) | 14 | 20-32 | RSVP / Oddball paradigm | Single-short oddball trials > Search no-target trials | 3T | *Event-related* | N | N | 240 mm | Whole-brain | 7 | Bilateral | Brain Voyager QX 2.3 | Talairach |
| Han and Marois (2014) | 14 | 20-32 | RSVP / Oddball paradigm | Target > Distractor trials | 3T | *Event-related* | N | N | 240 mm | Whole-brain | 11 | Bilateral | Brain Voyager QX 2.3 | Talairach |
| Han and Marois (2014) | 6 | 19-35 | RSVP / Oddball paradigm | Target > Distractor trials | 3T | *Event-related* | N | N | 240 mm | Whole-brain | 11 | Bilateral | Brain Voyager QX 2.3 | Talairach |
| Han et al. (2018) | 20 | 22-33 | RSVP / Oddball paradigm | Oddball > Search trials | 3T | *Event-related* | N | N | 240 mm | Whole-brain | 6 | Bilateral | FSL | MNI |
| Harding et al. (2016) | 25 | 25.5 ± 4.4 | Working memory paradigm | 2-back > 0-back trials (main effect) | 3T | *Event-related* | N | N | 210 x 210 mm | Whole-brain | 21 | Bilateral | SPM8 | MNI |
| Henseler et al. (2011) | 27 | 24.56 ± 2.53 | Working memory paradigm | External attending to targets > Baseline | 3T | Blocked | N | N | 192 mm | Whole-brain | 7 | NA | SPM2 | MNI |
| Henseler et al. (2011) | 27 | 24.56 ± 2.53 | Working memory paradigm | Internal orienting > Baseline | 3T | Blocked | N | N | 192 mm | Whole-brain | 8 | NA | SPM2 | MNI |
| Henseler et al. (2011) | 27 | 24.56 ± 2.53 | Working memory paradigm | Internal > External attending (position task) | 3T | Blocked | N | N | 192 mm | Whole-brain | 15 | Bilateral | SPM2 | MNI |
| Kim et al. (2012) | 16 | 23.6 ± 2.9; 18-35 | Task-switching paradigm | Task switch > Non-switch trials (main effect) | 3T | *Event-related* | N | N | 224 mm | Whole-brain | 20 | Left | SPM5 | MNI |
| Kim et al. (2012) | 16 | 23.6 ± 2.9; 18-35 | Stroop paradigm | Incongruent > Congruent trials (main effect) | 3T | *Event-related* | N | N | 224 mm | Whole-brain | 22 | Bilateral | SPM5 | MNI |
| Lin et al. (2019) | 18 | 27.4 ± 6.6 | Working memory paradigm | FC face cue and target > FC scene cue and target | 3T | *Event-related* | Y | N | 220 mm | Whole-brain | 9 | Right | FSL | MNI |
| Melcher and Gruber (2006) | 12 | 25.67 ± 1.88 | Stroop paradigm | Incongruent > Congruent trials | 3T | *Event-related* | N | N | 192 mm | Whole-brain | 13 | NA | SPM2 | MNI |
| Melcher and Gruber (2006) | 12 | 25.67 ± 1.88 | Oddball paradigm | Word-oddball vs. Oddball control | 3T | *Event-related* | N | N | 192 mm | Whole-brain | 19 | Bilateral | SPM2 | MNI |
| Melcher and Gruber (2006) | 12 | 25.67 ± 1.88 | Oddball paradigm | Color-oddball vs. Oddball control | 3T | *Event-related* | N | N | 192 mm | Whole-brain | 29 | Bilateral | SPM2 | MNI |
| Roth et al. (2006) | 12 | 23; 19-34 | Working memory paradigm | Sustanined working memory > Sustained feature detection | 3T | Blocked | N | N | N/A | Whole-brain | 15 | Left | AFNI | Talairach |
| Roth et al. (2006) | 12 | 23; 19-34 | Working memory paradigm | Update > Cued maintenance | 3T | *Event-related* | N | N | N/A | Whole-brain | 11 | Left | AFNI | Talairach |
| Sreenivasan et al. (2014) | 16 | 22; 18-32 | Working memory paradigm | Memory > Rotation discrimination trials | 3T | Blocked | N | N | N/A | ROIs | 13 | Right | AFNI | Talairach |
| Stelzel et al. (2011) | 48 | F: 22 ± 1.99; M: 22.6 ± 1.99 | Task-switching paradigm | Task switch > Repetition trials | 3T | *Event-related* | N | N | 240 mm | Whole-brain | 5 | Left | SPM5 | MNI |
| Todd et al. (2011) | 18 | 18-31 | Working memory paradigm | Encoding period > Baseline | 3T | *Event-related* | N | N | 240 mm | Whole-brain | 27 | Bilateral | BrainVoyager QX v.1.09 | Talairach |
| Wills et al. (2017) | 22 | 18-30 | Contingent capture paradigm | Salient target > Baseline | 3T | *Event-related* | N | N | 220 mm | ROIs | 13 | Bilateral | AFNI | Talairach |
| Yin et al. (2017) | 26 | 21.3; 21-25 | Task-switching paradigm | Task switch > Repetition trials | 3T | *Event-related* | N | N | 192 mm | Whole-brain | 15 | Left | SPM8 | MNI |
| Zanto et al. (2010) | 13 | 25; 20-31 | Working memory paradigm | FC with V4: Attend > Ignore color | 3T | *Event-related* | N | Y | 230 mm | Whole-brain | 5 | Right | SPM5 | MNI |
| Zanto et al. (2010) | 13 | 25; 20-31 | Working memory paradigm | FC with V5: Attend > Ignore motion | 3T | *Event-related* | N | Y | 230 mm | Whole-brain | 8 | Bilateral | SPM5 | MNI |
| Zanto et al. (2011) | 20 | 24.25; 18-31 | Working memory paradigm | FC with V4: Attend > Ignore color | 3T | *Event-related* | N | Y | 230 mm | Whole-brain | 18 | Right | SPM5 | MNI |
| Zanto et al. (2011) | 20 | 24.25; 18-31 | Working memory paradigm | FC with V5: Attend > Ignore motion | 3T | *Event-related* | N | Y | 230 mm | Whole-brain | 16 | Bilateral | SPM5 | MNI |
| Zhang et al. (2018) | 19 | 19-26 | Feature-based attention paradigm | Stimulus block > Baseline | 3T | Blocked | Y | N | 192 mm | ROIs | 8 | Bilateral | BrainVoyager QX | Talairach |
| Zhao et al. (2020) | 13 | 21.54 ± 1.99 | Working memory paradigm | High working memory load > Low working memory load | 3T | *Event-related* | N | N | 210 mm | Whole-brain | 7 | Bilateral | SPM8 | MNI |

### 1. ALE control and contrast analyses method

We carried out two control analyses in the FEF and IFJ localizer samples to rule out potential biases in our main results. As we were mainly interested in inferring the localization of the FEF and IFJ, we decided to include studies with partial brain coverage. We motivated this choice by accepting the tradeoff between having access to a larger sample of experiments for these regions, as opposed to having less sensitivity in detecting other regions that are consistently active during the tasks included (which however are not the main focus of the present study), but not reported simply due to the lack of whole-brain brain coverage. The inclusion of studies with partial brain coverage is however generally not recommended in ALE analyses (Müller et al. 2018). In our first set of control analyses, we have therefore excluded those studies based on the FOV parameters reported in the study (see Tables S1 and S2) or the author's description of the fMRI data acquisition. This led to the exclusion of two experiments from the FEF sample (namely Amiez and Petrides 2018, and Berman et al. 1999), and one study in the IFJ sample (Sreenivasan et al. 2014). For this control analysis, the ALE parameters were set to 5000 threshold permutations and a voxel-level FWE of 0.01 with a minimum cluster size of $50mm^3$ (corresponding to 6 voxels). The second set of control analyses was carried out to rule out another potential source of biases, that is, ROI analyses (Müller et al. 2018). These analyses imply restricting the assessment of statistical significance by some form of spatial masking. In this case, we excluded all the studies that either carried out ROI analyses or that only reported their results for specific ROIs (or even only FEF and IFJ foci; see Tables S1 and S2, column "Results reported for"). This led to an important decrease in both the FEF and IFJ localizer sample sizes (to 16 and 22 eligible experiments, respectively). Again, the ALE parameters were set to 5000 threshold permutations and a voxel-level FWE of 0.01 with a minimum cluster size of $50mm^3$ (corresponding to 6 voxels), as in our main localizer analyses.

As we were also interested in exploring potential dissociations in the prefrontal cortex, we carried out several ALE contrast analyses. For all these contrast analyses, we first computed separate ALE results for each function or contrast of interest by setting the ALE parameters to 10000 threshold permutations and a cluster-level FWE of 0.01 with $p < 0.001$. The experiments included in either analysis were randomly split into two groups of equal size to run a third pooled ALE analysis using the same parameters. The ALE scores of these two groups were then subtracted voxel-wise from each other 10000 times to create a null distribution of the difference between them. We used a threshold of $p < 0.01$ to infer significant differences between the two groups with a minimum cluster size of 25 $mm^3$ (as in Cieslik et al. 2016). The voxel-wise minimum of the ALE scores between the samples was used to create the conjunction image that indicates the similarity between the two groups of experiments.

### 2. ALE control and contrast analyses - FEF sample

1. **Cluster-level FWE control analysis.** In our main FEF localizer analysis, we applied voxel-level FWE correction, as this method allowed us a fine-grained assessment of the significant activations near the FEF. However, in the case of regions that are under-reported in the literature (i.e., the iFEF; Derrfuss et al. 2012), the use of a voxel-level FWE correction method may impose a too-conservative threshold that would prevent us from detecting clusters of activity that are also consistently activated in the FEF localizer sample. For instance, this could be due to the fact that most studies where an FEF functional localizer was employed tend to report only a pair of bilateral foci, leaving out other regions potentially active in the task (see Table S1 for the studies that have only reported part of their results), thus biasing the results in favor of the main activation cluster. In this first control analysis, we applied a less stringent multiple comparison correction method in order to reveal other consistently active peaks in neighboring frontal sites, namely cluster-level FWE. This correction method offers an optimal balance between sensitivity and specificity in ALE analyses to detect regions that may be under-reported (Eickhoff et al. 2016). As we introduced above, since the iFEF is under-reported in the fMRI literature, we hypothesized that this control analysis could be helpful in uncovering these activations. We therefore repeated the same ALE procedure on the FEF localizer sample setting the ALE parameters to a p-value of 0.001, with 10000 threshold permutations and applying a cluster-level FWE of 0.01.

2. **ALE contrast analysis - antisaccades vs prosaccades.** The second control analysis was intended as a replication of the ALE meta-analysis by Cieslik et al. (2016), who performed an ALE contrast analysis between experiments investigating prosaccades > fixation vs antisaccades > prosaccades in two equal samples of 12 experiments (see also Jamadar et al. 2013, for previous results). We note that, unlike Cieslik et al. (2016), we did not include PET studies, and also 1.5T fMRI studies to increase the spatial accuracy of our analysis (see the Supplementary Table 4 below for the details of the experiments included).

3. **ALE contrast analysis - prosaccades vs covert spatial attention.** In our main localizer analysis, we did not include covert spatial attention tasks as these aren't considered the gold standard to localize the FEF. However, covert spatial attention paradigms are often employed to localize all the main nodes of the dorsal attention network, including the FEF (Corbetta and Shulman 2002). Therefore, studies using adaptations of the spatial cueing paradigm (Posner et al. 1980) were included in this analysis (see Supplementary Table 5). Running a contrast analysis between prosaccades and covert spatial attention paradigm allowed us to compare the topography and the sources of overt and covert attention near the putative FEF. For this analysis, we used prosaccades > fixation vs valid > neutral/invalid trials contrasts.

#### 3. ALE contrast analyses - IFJ sample

Given the heterogeneity of the experiments included in the IFJ localizer sample, we carried out exploratory ALE contrast analyses by splitting up the IFJ sample according to the function investigated by each paradigm (i.e., oddball/attention vs working memory vs task-switching and Stroop paradigms; see Table S2) to see whether these paradigms elicited activity in distinct regions near the putative IFJ and to assess potential lateralization patterns. Task-switching and Stroop paradigms were grouped based on the results from Derrfuss et al. (2005), and we refer to this group of experiments as targeting cognitive control. In the first group of experiments (oddball/attention), most of the oddball experiments contrasted oddball > target trials and target trials > baseline (see for example Asplund et al. 2010), thus tapping on both stimulus-driven and goal-driven attentional mechanisms. There were also some examples of blocked design cueing paradigms contrasting target vs fixation blocks (Baldauf and Desimone 2014; Zhang et al. 2018). In the working memory group, the majority of experiments contrasted the functional connectivity between perceptual seed regions (V4, V5, FFA) in the attended vs ignored conditions (see for example Bollinger et al. 2010; Lin et al. 2019; Zanto et al. 2010, 2011). We note that even though these studies were based on a second-level contrasts analysis, and thus involve some form of masking (i.e., the restriction to a specific brain region for the assessment of the significance level, hence spatial bias), the significant correlations with each seed were assessed over the whole-brain, making the localization of the IFJ with this method likely only slightly affected by this issue, if at all. Indeed, as previously suggested in Nee et al. (2013), their results match well those from traditional univariate analyses. The cognitive control group was mostly composed of experiments contrasting switch > repeat trials and incongruent > congruent contrasts in Stroop paradigms.

We therefore ran three separate ALE contrast analyses:

1. **Oddball/attention vs working memory** (11 vs 12 experiments):
2. **Working memory vs cognitive control** (12 vs 11 experiments), and;
3. **Oddball/attention vs cognitive control** (11 vs 11 experiments).

**Supplementary Table 3** List of the experiments included in the prosaccade > fixation analysis

| Study | N | Age | Paradigm | Contrast | Scanner | Design | Eye tracker | Results reported for | N° of foci | Multiple FEF foci | Software | Space |
|---|---|---|---|---|---|---|---|---|---|---|---|---|
| Alkan et al. (2011) | 8 | 26 ± 4 | Functional localizer | Prosaccades > Fixation | 3T | Blocked | Y | Whole-brain | 19 | N | AFNI | Talairach |
| Amiez and Petrides (2018) | 13 | 22.6 ± 2.8 | Functional localizer | Prosaccades > Fixation | 3T | Blocked | N | FEF foci | 2 | N | SPM12b | MNI |
| Atmaca et al. (2013) | 11 | 26.9; 22-33 | Functional localizer | Prosaccades > Fixation | 3T | Blocked | N | Whole-brain | 24 | Y | SPM8 | MNI |
| Bär et al. (2016) | 14 | 22-56 | Functional localizer | Prosaccades > Fixation | 3T | Blocked | N | Whole-brain | 12 | N | BrainVoyager QX 1.10 | Talairach |
| Berman et al. (1999) | 11 | 25.6 ± 7.1; 18-43 | Functional localizer | Prosaccades > Fixation | 3T | Blocked | N | ROIs | 14 | Y | AFNI | Talairach |
| Braga et al. (2016) | 20 | 26.2; 21-36 | Functional localizer | Prosaccades > Fixation | 3T | Blocked | Y | Whole-brain | 7 | N | FSL | MNI |
| Brown et al. (2006) | 10 | 26; 22-33 | Functional localizer | Prosaccades > Fixation | 4T | *Event-related* | Y | Whole-brain | 17 | N | BrainVoyager 2000 | Talairach |
| Connolly et al. (2000) | 7 | 24.8 ± 3.2 | Functional localizer | Prosaccades > Fixation | 4T | Blocked | N | Whole-brain | 7 | Y | Stimulate | Talairach |
| Connolly et al. (2002) | 8 | NA | Functional localizer | Prosaccades > Fixation | 4T | Blocked | N | ROIs | 4 | N | Stimulate | Talairach |
| Connolly et al. (2005) | 5 | NA | Functional localizer | Prosaccades > Fixation | 4T | Blocked | Y | ROIs | 5 | N | Stimulate / Brain Voyager 4.9 | Talairach |
| Connolly et al. (2007) | 8 | NA | Functional localizer | Prosaccades > Fixation | 4T | Blocked | N | ROIs | 2 | Y | Stimulate / BrainVoyager 4.9 / QX | Talairach |
| Christophel et al. (2018) | 22 | 24.4 ± 0.83 | Functional localizer | Prosaccades > Fixation | 3T | *Event-related* | N | FEF foci | 2 | N | SPM8 | MNI |
| Duecker et al. (2013) | 20 | 19-28 | Functional localizer | Prosaccades > Fixation | 3T | Blocked | N | FEF foci | 2 | N | BrainVoyager QX 2.3 | Talairach |
| Frannson et al. (2014) | 24 | 26.7 ± 4.9 | Functional localizer | Prosaccades > Fixation | 3T | Blocked | Y | Whole-brain | 8 | N | SPM8 | MNI |
| Guo et al. (2012) | 12 | 19-31 | Functional localizer | Prosaccades > Fixation | 3T | Blocked | Y | ROIs | 6 | N | SPM8 | Talairach |
| Gurel et al. (2018) | 16 | 23.2 | Functional localizer | Prosaccades > Fixation | 3T | Blocked | Y | FEF foci | 2 | N | BrainVoyager QX | Talairach |
| Heinen et al. (2006) | 5 | 20-37 | Functional localizer | Prosaccades > Fixation | 3T | Blocked | Y | ROIs | 4 | N | VISTASOFT | Talairach |
| Hubl et al. (2008) | 7 | 31 ± 9 | Functional localizer | Prosaccades > Fixation | 3T | Blocked | N | Whole-brain | 9 | N | BrainVoyager QX | Talairach |
| Jamadar et al. (2015) | 23 | 25.8; 18-43 | Functional localizer | Prosaccades > Fixation | 3T | *Event-related* | Y | Whole-brain | 47 | N | SPM8 | MNI |
| Jarvstad & Gilchrist (2019) | 23 | NA | Functional localizer | Prosaccades > Fixation | 3T | Blocked | Y | Whole-brain | 4 | N | FSL v.5.06-1 | MNI |
| Kastner et al. (2007) | 4 | 20-36 | Functional localizer | Prosaccades > Fixation | 3T | Blocked | Y | ROIs | 5 | N | AFNI | Talairach |
| Kurata et al. (2005) | 6 | 29-38 | Functional localizer | Prosaccades > Fixation | 3T | Blocked | N | Whole-brain | 14 | N | Advanced Visual Systems & AFNI | Talairach |
| Levy et al. (2007) | 4 | 24-43 | Functional localizer | Prosaccades > Fixation | 3T | Blocked | N | ROIs | 5 | N | BrainVoyager QX | Talairach |
| Matsuo et al. (2003) | 12 | 20-42 | Functional localizer | Prosaccades > Fixation | 3T | Blocked | N | FEF foci | 1 | N | SPM99 | MNI |
| Pierce et al. (2019) | 30 | 25.7 ± 4.1 | Functional localizer | Prosaccades > Fixation | 3T | Blocked | Y | Whole-brain | 9 | Y | SPM12 | MNI |
| Schon et al. (2008) | 17 | 21.29 ± 3.72 | Functional localizer | Prosaccades > Fixation | 3T | Blocked | N | Whole-brain | 19 | N | SPM2 | MNI |
| Tamber-Rosenau et al. (2018) | 8 | 28.5 ± 3.3 | Functional localizer | Prosaccades > Fixation | 3T | Blocked | Y | ROIs | 6 | N | BrainVoyager QX v.1.10.2–2.8 | Talairach |
| Tark & Curtis (2009) | 5 | 22-39 | Functional localizer | Prosaccades > Fixation | 3T | Blocked | Y | FEF foci | 2 | N | Caret | MNI |
| Tibber et al. (2010) | 16 | 21-40 | Functional localizer | Prosaccades > Fixation | 3T | Blocked | Y | ROIs | 9 | N | SPM5 | Talairach |

**Supplementary Table 4** List of the experiments included in the antisaccades > prosaccade analysis

| Study | N | Age | Paradigm | Contrast | Scanner | Design | Eye tracker | Results reported for | N° of foci | Software | Space |
|---|---|---|---|---|---|---|---|---|---|---|---|
| Brown et al. (2006) | 10 | 26; 22-33 | Antisaccade paradigm | Antisaccades > Prosaccades trials | 4T | *Event-related* | Y | Whole-brain | 15 | BrainVoyager 2000 | Talairach |
| Brown et al. (2007) | 11 | 25; 20-28 | Antisaccade paradigm | Antisaccades > Prosaccades trials | 4T | *Event-related* | Y | Whole-brain | 14 | BrainVoyager 2000 | Talairach |
| Brown et al. (2007) | 11 | 25; 20-28 | Antisaccade paradigm | Antisaccades > Prosaccades trials | 4T | *Event-related* | Y | Whole-brain | 11 | BrainVoyager 2000 | Talairach |
| Cameron et al. (2009) | 11 | 22-30 | Antisaccade paradigm | Antisaccades > Prosaccades trials | 3T | *Event-related* | Y | FEF foci | 2 | BrainVoyager 1.9 | Talairach |
| Ford et al. (2005) | 10 | 28 | Antisaccade paradigm | Antisaccades > Prosaccades trials | 4T | *Event-related* | Y | Whole-brain | 8 | BrainVoyager 2000 v. 4.8 | Talairach |
| Manoach et al. (2007) | 21 | 34.2 ± 12.6 | Antisaccade paradigm | Antisaccades > Prosaccades trials | 3T | *Event-related* | Y | Whole-brain | 7 | FreeSurfer | Talairach |
| Neggers et al. (2012) | 13 | 20-35 | Antisaccade paradigm | Antisaccades > Prosaccades trials | 3T | *Event-related* | N | ROIs | 13 | SPM5 | MNI |
| Peirce & McDowell (2016) | 35 | 19 ± 3.5 | Antisaccade paradigm | Antisaccades > Prosaccades trials | 3T | *Event-related* | Y | Whole-brain | 5 | AFNI | Talairach |
| Peirce & McDowell (2017) | 35 | 19 ± 3.5 | Antisaccade paradigm | Antisaccades > Prosaccades blocks | 3T | Blocked | Y | Whole-brain | 2 | AFNI | Talairach |
| Salvia et al. (2020) | 14 | 27.1 ± 2.7 | Antisaccade paradigm | Antisaccades > Prosaccades trials | 3T | *Event-related* | Y | Whole-brain | 12 | SPM12 | MNI |

**Supplementary Table 5** List of the experiments included in the endogenous covert spatial attention analysis

| Study | N | Age | Paradigm | Contrast | Scanner | Design | Eye tracker | Results reported for | N° of foci | Multiple FEF foci | Software | Space |
|---|---|---|---|---|---|---|---|---|---|---|---|---|
| Chica et al. (2013) | 18 | 25 ± 4 | Spatial cueing | Cue > Jitter Fixation | 3T | *Event-related* | N | Whole-brain | 22 | Y | SPM5 | MNI |
| Fan et al. (2005) | 16 | 27.2 ± 5.7 | Attentional network test | Spatial > Center cue | 3T | *Event-related* | N | Whole-brain | 8 | N | SPM99 | MNI |
| Ikkai & Curtis (2008) | 14 | 21-35 | Attentional cueing | Covert shift > Baseline | 3T | *Event-related* | Y | ROIs | 11 | Y | Caret | MNI |
| Mao et al. (2007) | 12 | 24.3; 21-27 | Spatial cueing | Attention to peripheral locations > Fixation | 3T | Blocked | N | Whole-brain | 6 | N | SPM99 | Talairach |
| Mohanty et al. (2009) | 13 | 27 ± 3.8 | Spatial cueing / Visual search | Valid > Neutral trials | 3T | *Event-related* | Y | Whole-brain | 10 | N | SPM5 | MNI |
| Vossel et al. (2012) | 24 | 26.83; 20-37 | Spatial cueing | Valid trials > Baseline | 3T | *Event-related* | Y | Whole-brain | 6 | N | SPM8 | MNI |
| Wen et al. (2012) | 12 | 20-28 | Spatial cueing | Attend > Passive viewing blocks | 3T | Blocked | N | ROIs | 7 | N | SPM2 | MNI |
| Xuan et al. (2016) | 24 | 26.3; 18-49 | Attentional network test | Valid > Invalid trials | 3T | *Event-related* | N | Whole-brain | 20 | N | SPM8 | MNI |

#### 4. ALE control analyses results

The results of the first control analysis, in which we excluded studies with partial brain coverage, replicated our main localizer results both in terms of the inferred ALE peaks and the number and anatomical localization of the significant clusters. In the FEF sample, the ALE peaks coincided with the main analysis, and the only difference that emerged was a minor shift of the left iFEF peak (from MNI152: -48, -2, 40 to -46, -2, 38). In the IFJ sample, the ALE peaks again coincided with the results of our main analysis.

The results of the second control analysis where we excluded all ROI analyses again matched well with our main results in terms of the inferred ALE peaks but had important differences regarding the number and anatomical localization of the significant clusters. The results of this control analysis are reported in Table S6 and Table S7 for the FEF and IFJ samples, respectively.

In the FEF sample, the cluster with the highest ALE value was now localized primarily in the left medial frontal gyrus, including the SCEF. The second and third clusters larger clusters were however again localized in the left and right FEF respectively and were approximately of the same size. In terms of the inferred FEF ALE peaks, these were almost identical to our main results (LH main: -28, -6, 54 vs no ROI control: -28, -6, 56; and RH main: 30, -6, 50 vs no ROI control: 28, -6, 50; all the coordinates are in MNI152 space). However, in contrast to our main analysis results, the left FEF cluster did not spread onto the iPCS and there wasn't any peak corresponding to the iFEF. Interestingly, we also found a second peak in the right FEF cluster, which was lateral relative to the first, although still localized within the sPCS (MNI152: 40, -4, 50). Similar to the main results, there were other two clusters localized in the left and right precuneus and SPL, but we didn't find evidence of a cluster in the right IPL, possibly due to the lower sample size (35  vs 16 experiments).

In the IFJ sample, the larger clusters were localized in the left and right IFJ, replicating our main analysis results. Their ALE peak exactly matched those from our main results in the left hemisphere (-42, 6, 30) but had some slight differences in the right hemispheres, where we found two peaks with similar ALE values (main; 46, 12, 28 vs no ROI control: 46, 10, 26, and 42, 8, 30; all the coordinates are in MNI152 space). This indicates some residual variability in the localization of the right IFJ, which wasn't detectable in our main analysis results. The remaining clusters were similar to the main analysis results, with the exception that the cluster in the left SPL/IPL was now disjointed into two smaller clusters.

**Supplementary Table 6** FEF sample no ROI control analysis ALE results

| Cluster | Macroanatomical location | Hemi | MNI152 coordinates | | | ALE value | Volume (mm³) | BA |
|---|---|---|---|---|---|---|---|---|
| | | | x | y | z | | | |
| 1 | Medial Frontal Gyrus | L/R | -2 | 0 | 58 | 0.0386 | 1360 | 6 |
| 2 | Precentral Gyrus | L | -28 | -6 | 56 | 0.0341 | 992 | 6 |
| 3 | Middle Frontal Gyrus | R | 28 | -6 | 50 | 0.0308 | 960 | 6 |
| | Precentral Gyrus | R | 40 | -4 | 50 | 0.0275 | | 6 |
| 4 | Precuneus | L | -22 | -58 | 56 | 0.0306 | 488 | 7 |
| 5 | Precuneus | R | 26 | -60 | 56 | 0.0223 | 112 | 7 |

**Supplementary Table 7** IFJ sample no ROI control analysis ALE results

| Cluster | Macroanatomical location | Hemi | MNI152 coordinates | | | ALE value | Volume (mm³) | BA |
|---|---|---|---|---|---|---|---|---|
| | | | x | y | z | | | |
| 1 | Precentral Gyrus | L | -42 | 6 | 30 | 0.0532 | 2632 | 6 |
| 2 | Inferior Frontal Gyrus | R | 46 | 10 | 26 | 0.0402 | 1672 | 9 |
| | Precentral Gyrus | R | 42 | 8 | 30 | 0.0389 | | 6 |
| 3 | Medial Frontal Gyrus | L | -2 | 18 | 46 | 0.0462 | 1656 | 6 |
| 4 | Precentral Gyrus | L | -28 | -4 | 54 | 0.0474 | 816 | 6 |
| 5 | Claustrum | R | 32 | 22 | -4 | 0.0348 | 456 | * |
| 6 | Precuneus | L | -26 | -66 | 44 | 0.0302 | 456 | 7 |
| 7 | Inferior Parietal Lobule | L | -32 | -54 | 48 | 0.0348 | 384 | 7 |
| 8 | Superior Parietal Lobule | R | 34 | -54 | 46 | 0.0323 | 312 | 7 |
| 9 | Claustrum | L | -30 | 18 | 2 | 0.0317 | 264 | * |

#### 5. ALE control and contrast analyses - FEF sample results

By repeating the ALE analysis using cluster-level FWE instead of voxel-level FWE, we were able to uncover bilateral activations ventral to the main FEF ALE peaks. These activations were extending from the sPCS to the posterior bank of the iPCS and were primarily localized in the iPCS (see Figure S3), corresponding to the iFEF (Kastner et al. 2007). This highlights the fact that, although these clusters may be under-reported in the literature (Derrfuss et al. 2012), they were nevertheless consistently activated in our sample of experiments.

Our two ALE contrast analyses show evidence of important spatial segregation and overlap within FEF for antisaccades, prosaccades and covert spatial attention contrasts (see Figures S4 and S5). Starting with the first contrast analysis, we would like to comment separately on the ALE results for antisaccades > prosaccades and the 'pure' prosaccade > fixation results, as these give important clues on how to interpret the ALE peaks from our main analysis results (where we also included 'mixed' antisaccades & prosaccades > fixation and antisaccades > fixation contrasts to increase our sample size). In the prosaccades > fixation ALE results the FEF peaks were localized in: LH 1st ALE peak (-28, -6, 54); LH 2nd ALE peak (-52, 0, 40); RH 1st ALE peak (36, -4, 52); RH 2nd ALE peak (30, -4, 50); RH 3rd ALE peak (34, -2, 50). While the left hemisphere peaks match the main analysis results, in the right hemisphere the strongest convergence is now found more lateral compared to the main result ALE peak (30, -6, 50) suggesting a higher variability along this axis (see Figure S4, Panel A). In the antisaccades > prosaccades ALE results there were only two peaks localized within FEF, specifically in LH (-26, -2, 56), and RH (-28, -2, 54). These peaks were localized more anteriorly and medially compared to our main results (Figure S4, Panel B). This already suggests that there may be important dissociations between the activations derived from these contrasts. Indeed, when we directly contrasted these two samples, we found that the antisaccades > prosaccades and prosaccades > fixation experiments consistently activated segregated prefrontal clusters, as previously reported by other studies (Cieslik et al. 2016; Jamadar et al. 2013). While in the right hemisphere, the antisaccade cluster was medial relative to the prosaccade cluster, and largely overlapping at the junction of the sPCS with the SFS, this organization was less evident in the left hemisphere, where the two clusters overlapped near the same anatomical location, but with a segregated cluster for antisaccades localized in its anterior-medial part (see Figure S4, Panel C). Overall, these results suggest that the additional processes required for the antisaccade task (Munoz and Everling 2004), namely response inhibition and the execution of a saccade towards the opposite location relative to the target, consistently recruit the medial part of the FEF (McDowell et al. 2008). These processes may be mediated by distinct neural populations within the FEF (see Lowe and Schall 2018, for a classification in the macaque). We also found two bilateral segregated clusters in the medial prefrontal cortex for prosaccades localized in the posterior SCEF. The antisaccades clusters were instead localized in the right anterior SCEF. We did not find clusters in the posterior parietal cortex as in Cieslik et al. (2016), with the exception of a cluster in the left precuneus/SPL, possibly due to the fact that our antisaccade sample was lower compared to that study (8 vs 12 experiments), and the influence of studies only reporting ROI analyses.

Our second ALE contrast analysis, namely between prosaccades and covert spatial attention activations (Figure S5, Panel C) reveals a clear pattern of overlap near the junction of the sPCS and SFS. These results are consistent with the hypothesis that covert and overt attention have a spatially common source within the FEF (Astafiev et al. 2003; Corbetta et al. 1998; de Haan et al. 2008). We also found evidence of segregated clusters, with covert attention clusters being mostly localized in the posterior bank of the sPCS in the right and to a smaller extent in the left hemispheres (not visible in Figure S5). These results strongly suggest that covert spatial attention paradigms may be equally adept as FEF functional localizers compared to the current prosaccade gold standard, although more trials may be needed to reliably elicit activations in all subjects due to the weaker nature of the signal measured (Beauchamp et al. 2001; De Haan et al. 2008).

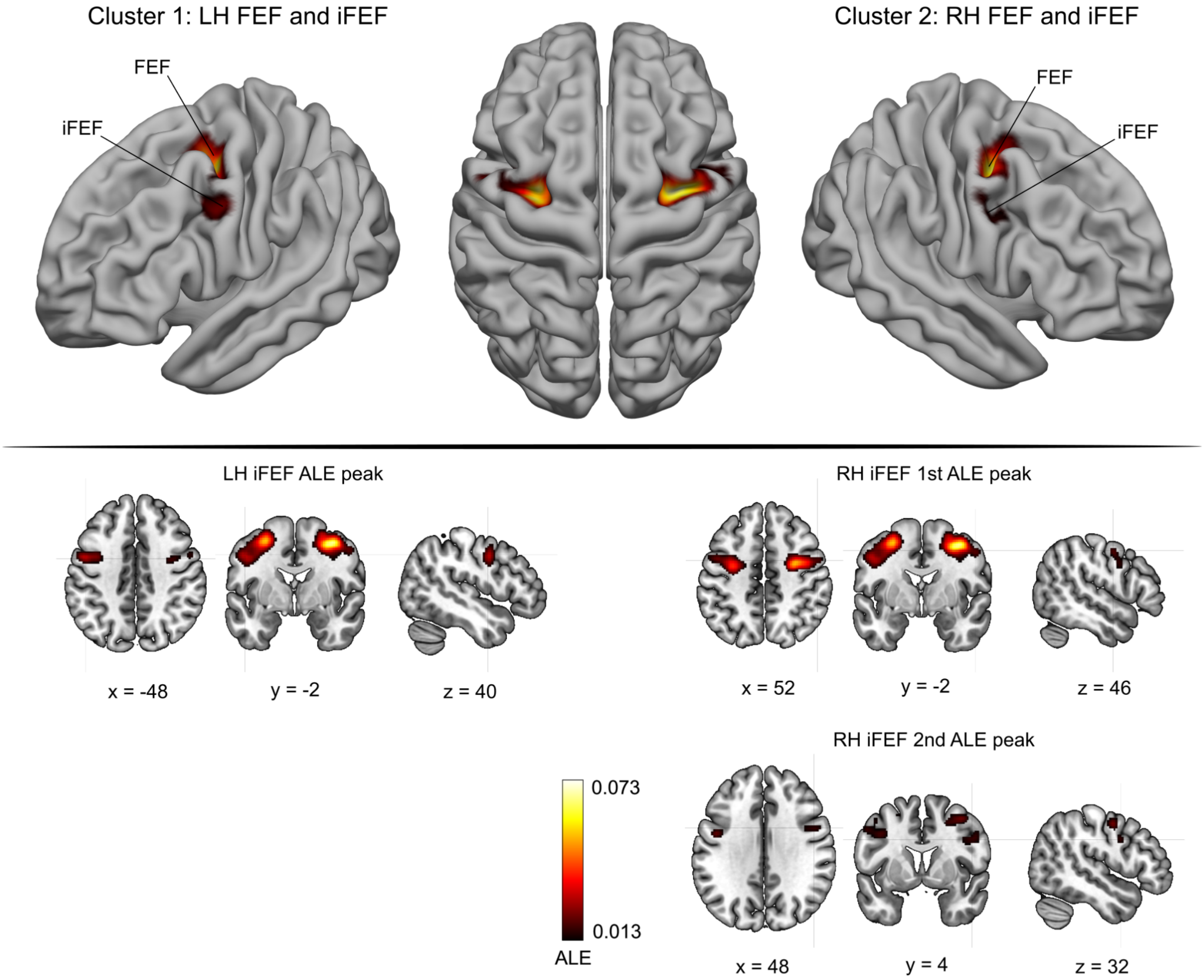


**Fig. S3** Results of the first FEF sample control analysis - main clusters of activation. Applying cluster-level FWE in the ALE analysis allowed us to uncover bilateral activations ventral to the main FEF peaks. These activations were extending from the sPCS to the posterior bank of the iPCS, and were primarily localized in the precentral gyrus. These results reveal the presence of consistent iFEF activations in the FEF localizer sample and three iFEF peaks (shown in the volumetric views of the figure)

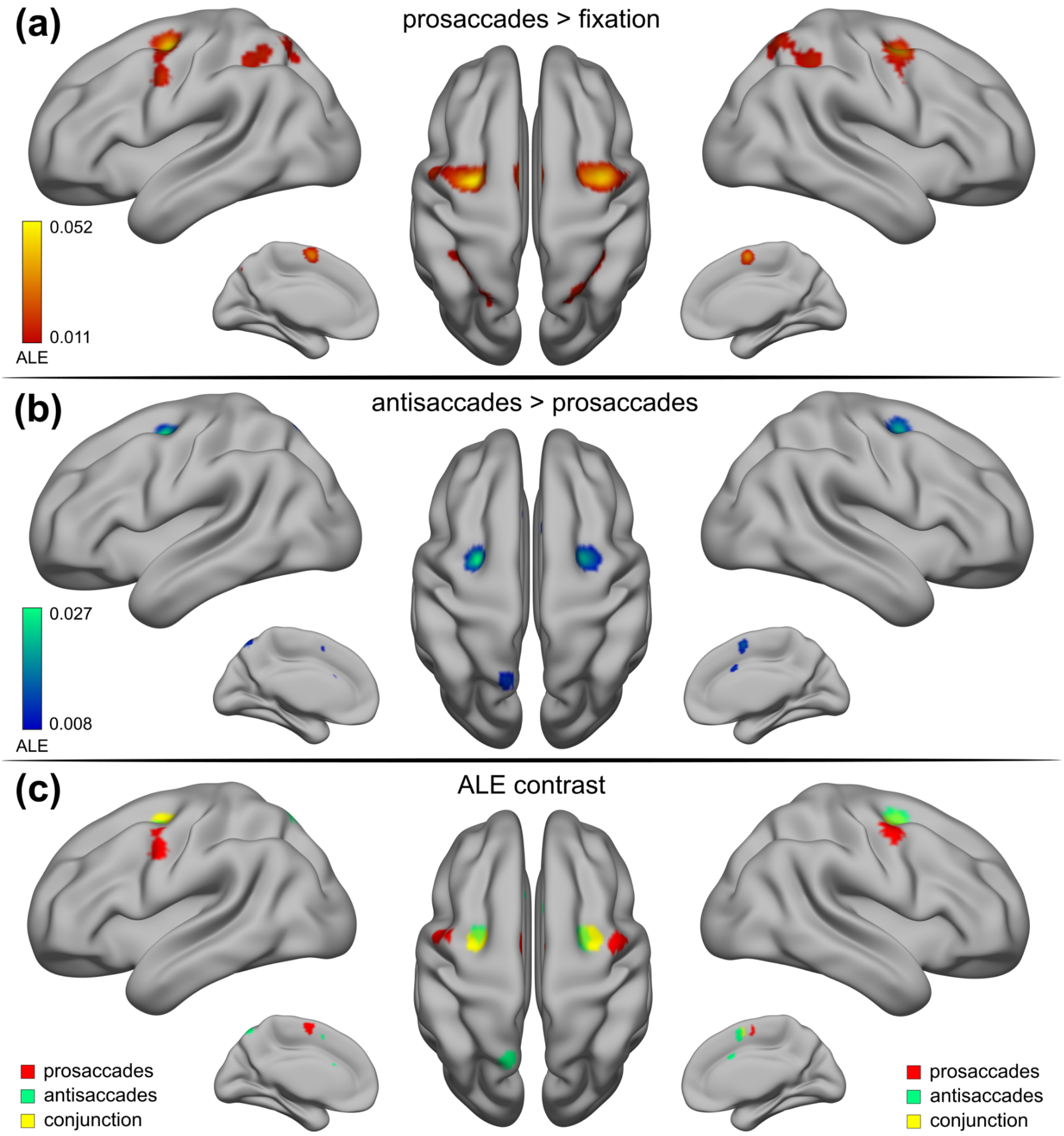


**Fig. S4** FEF sample contrast analysis between antisaccades > prosaccades vs prosaccades > fixation contrasts. **C** The clusters overlap in the bilateral posterior aspect of the SFS and sPCS. The contrast of these clusters showed that the antisaccades-related clusters were generally more medial and localized anterior to the prosaccades-related clusters. These results are consistent with those reported by Cieslik et al. (2016) and suggest that the additional mechanisms required for the antisaccade task are mediated by separate neural populations within the medial aspect of the FEF

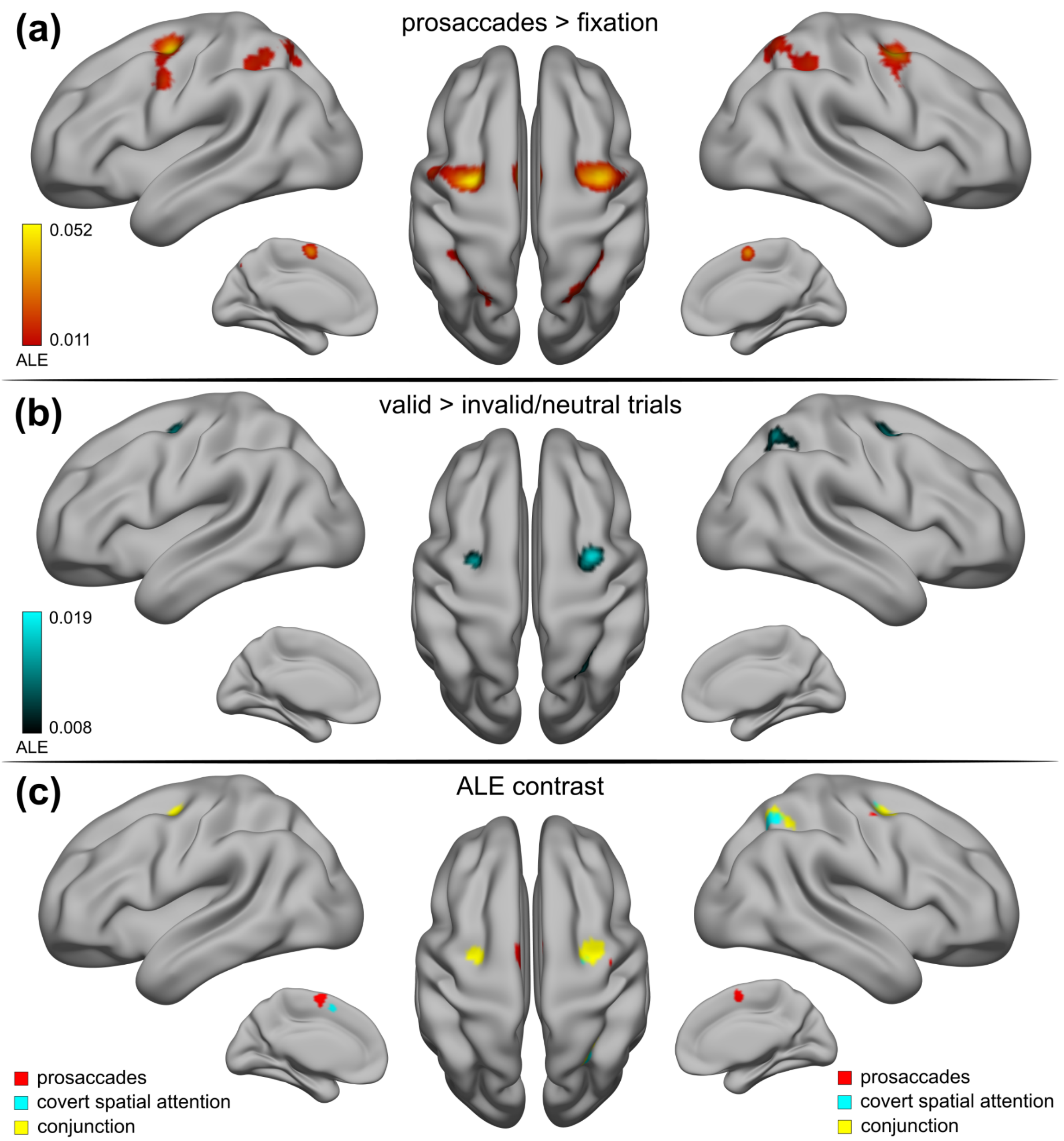


**Fig. S5** FEF sample contrast analysis between prosaccades > fixation vs valid > neutral/invalid trials in covert spatial attention task. **C** The clear overlap we found at the junction of the SFS and sPCS supports the hypothesis that covert and overt attention have a spatially common source within the FEF (Astafiev et al. 2003; Corbetta et al. 1998; de Haan et al. 2008)

#### 6. ALE contrast analyses - IFJ sample results

By splitting up our IFJ localizer sample based on the paradigm employed (i.e., oddball/attention vs working memory vs cognitive control), we found some dissociations within the putative IFJ, as well as interesting lateralization patterns (see Figure S6). While oddball/attention paradigms gave rise to bilateral activations near the putative IFJ, working memory paradigms consistently activated only a right hemisphere cluster, whereas in contrast cognitive control paradigms were only consistently activating on a cluster in the left hemisphere. Remarkably, each of these clusters had a quite distinct spatial topography. In the left hemisphere, the cluster related to cognitive control (task-switching and Stroop paradigms) extended from the posterior bank of the superior iPCS to the junction of the iPCS with the IFS, where it overlapped with the oddball/attention cluster (see Figure S6, Panel A). This cluster further extended anteriorly and ventrally. The same arrangement was approximately found in the right hemisphere, in which a posterior-dorsal working memory limited cluster overlapped with the oddball/attention cluster just above the junction of the iPCS with the IFS (see Figure S6, Panel B). The oddball/attention cluster again extended anteriorly and ventrally. Finally, the contrast between working memory and cognitive control only revealed overlap in the left paracentral and midcingulate cortex, as the other clusters were localized within the right and left hemispheres, respectively (see Figure S6, Panel C).

Based on the results from these ALE contrast analyses and their comparisons with our main results, we would like to offer some suggestions on how to effectively localize the IFJ and segregate it from adjacent brain regions. First of all, at the general level, contrasts involving two demanding experimental conditions (e.g., task switch > repeat trials) may be more appropriate to measure activity within the IFJ compared to experimental trials vs passive fixation conditions. Secondly, an even more stringent way to isolate this region would be to compare experimental conditions that are matched in difficulty, thus avoiding contamination from non-specific cognitive load effects (as in Baldauf and Desimone 2014). Oddball/attention paradigms seemed to tap on the same IFJ cluster as found in our main localizer (see Figure S6, Panel A). However, in the experiments we analyzed, these tasks usually contrast activity between oddball and target trials, leading to a low number of trials that are used as functional localizers within each run as a result (Han and Marois 2014). An additional problem may be caused by spatial smoothing, which would lead to merging activity with the IFG, a node classically viewed as belonging to the ventral attention network (Corbetta and Shulman 2002), which is also significantly activated in these paradigms (Levy and Wagner 2011). Therefore, a more straightforward way to isolate the IFJ from the IFG may be achieved by administering top-down feature- and object-based attention tasks (Baldauf and Desimone 2014; Liu et al. 2011; Liu 2016; Zhang et al. 2018) and contrasting valid > invalid trials collapsed across the stimuli features/dimensions (e.g., similarly to the approach reported in Zanto et al. 2010). Posteriorly, another important issue is how to segregate IFJ activity from the iFEF (or the PEF in the MMP1 taxonomy). Derrfuss et al. (2012) reported that by employing a Stroop paradigm and contrasting incongruent > congruent trials, they were able to isolate activity from the adjacent iFEF (which was activated by the execution of voluntary saccades in darkness) in the native space in all the subjects they analyzed. These results show a promising way to infer the posterior IFJ functional border. As suggested previously however in the case of the FEF results, the presence of significant voxels within the iFEF is particularly difficult to interpret given that, in the IFJ localizer sample, only two out of 32 experiments (see Table S2) employed strict monitoring of eye movements in the scanner. The possibility that the convergence in iFEF in this sample may be due to this confound cannot be therefore completely ruled out (Amiez and Petrides 2009; Kato and Miyauchi 2003). Therefore we suggest that future localization approaches aimed at separating IFJ and iFEF activations must ensure a robust way to prevent data contamination from oculomotor artifacts. Finally, as suggested in our Discussion, the combination of different tasks and the manipulation of task difficulty seems a promising approach to delineate the IFJ based on a conjunction analysis (as in Stiers and Goulas 2018).

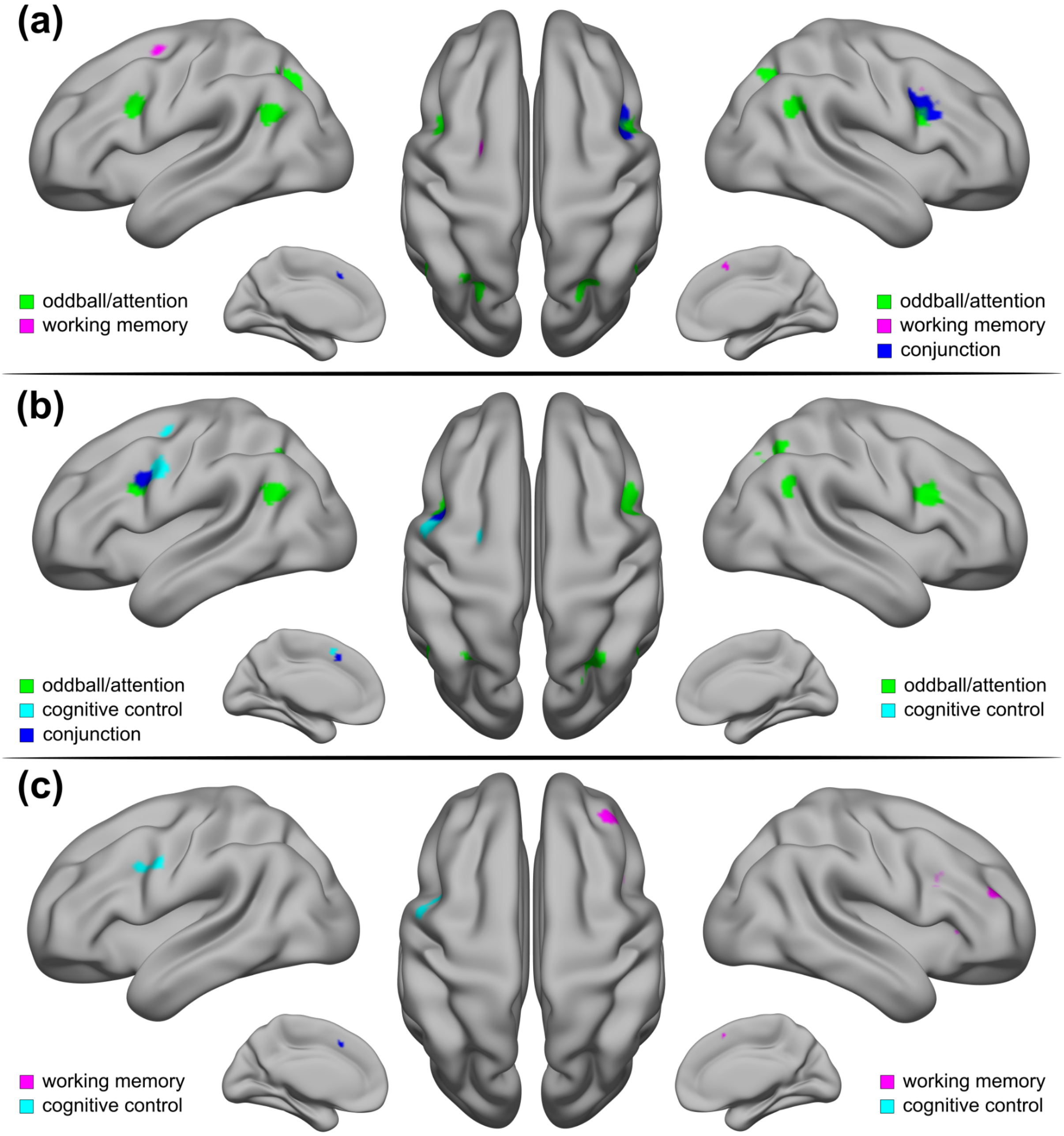


**Fig. S6** Results of the IFJ sample control analysis. The IFJ sample was split into three groups of experiments based on the paradigm employed (oddball/attention, working memory and cognitive control) to explore potential dissociations between them near the IFJ. In line with this possibility, we found a lateralized cluster involved in cognitive control (i.e., task-switching/Stroop paradigms) in the left hemisphere, and a cluster involved in working memory (mainly n-back contrasts) in the right hemisphere. These clusters overlapped with a bilateral cluster associated with oddball/attention experiments at the iPCS and IFS junction, including the IFJ. The oddball/attention clusters were however generally anterior and ventral relative to their location in both hemispheres (see **A** and **B**)

#### 7. MACM contrast analysis results

To quantitatively examine the dissociations in the MACM patterns of FEF and IFJ for each hemisphere, we also ran an ALE contrast analysis between the LH FEF vs LH IFJ, and RH FEF vs RH IFJ coactivation patterns. Our results generally confirm what can be seen in Figure 5. The LH IFJ coactivated with the RH IFJ and the LH FEF, whereas the LH FEF coactivated with the RH FEF, and LH and RH IFJ. The LH IFJ had also additional coactivations with the insular cortex. Medially, there was a posterior cluster in the SCEF differentially coactivated with the LH FEF, and two clusters in the anterior and mid cingulate cortex coactivated with the LH IFJ. Posteriorly, we found common activations in the IPL, but crucially, dissociations in the precuneus/SPL (for LH FEF) and the fusiform face complex, areas TE2p and V8, and the parahippocampal areas (for the LH IFJ). We found similar results in the right hemisphere. The RH FEF coactivated with the LH FEF, and the LH and RH IFJ. The RH IFJ only coactivated with the LH IFJ and had additional coactivations with the insular cortex. There were differential coactivations of the RH FEF with the precuneus/SPL and the lateral intraparietal area and common coactivations in the left SPL.

#### 8. MACM decoding results

Performing a reverse inference on the coactivation patterns of the LH FEF showed that the prevalent association was in the ‘action’ behavioral domain (see Figure 5, right side of panel A), namely execution.unspecified. In the ‘cognition’ domain, there were four prominent associations with attention, working memory, reasoning and spatial cognition. Finally, in the ‘perception’ domain, the two highest associations were with vision.motion and vision.shape. The behavioral domain associations with the coactivation patterns of the RH FEF (see Figure 5, right side of panel B) were very similar to the LH FEF. Again, the primary association in the ‘action’ domain was with execution.unspecified. The prevalent association was however in the ‘cognition’ domain with attention, followed by working memory, language.speech and by reasoning. As for the previous seed, the two highest associations were with vision.motion and vision.shape in the ‘perception’ domain. The functional decoding of the LH and RH IFJ coactivation patterns uncovered associations with similar behavioral domains, although with some interesting differences in their predominance. The LH IFJ coactivations had the highest association with attention in the ‘cognition’ domain (see Figure 5, right side of panel C), followed by language.semantics, working memory, and language.speech. The next strongest association was in the ‘emotion’ domain with positive.reward/gain. Then, there were significant associations in the ‘perception’ domain with vision.shape and vision.unspecified and audition. Finally, in the ‘action’ domain we found the most prevalent associations with inhibition and execution.unspecified. As for the previous seed, the RH IFJ had the highest association with attention in the ‘cognition’ domain (see Figure 5, right side of panel D). In the same domain, there were also strong associations with working memory, language.semantics, reasoning, and language.speech. Next, we found two prominent associations in the ‘action’ domain with inhibition and execution.unspecified. Again, we found an association with positive.reward/gain in the ‘emotion’ domain. Lastly, the strongest associations in the ‘perception’ domain were with vision.shape, audition and somesthesis.pain.

#### 9. Limitations

While we believe that the present study significantly advances our understanding of the localization and spatial organization of areas in the posterior-lateral PFC, we would like to acknowledge its limitations. First of all, even though the inclusion of ROI analyses is not recommended according to the best practices reported in Muller et al. (2018), we nevertheless decided to include them to increase our sample size, as we were mainly interested in inferring the localization of their peaks. When we excluded them in a control analysis (reported in the section 4 above), we showed that the inferred ALE remained virtually identical. We note however that the inclusion of these studies may have inflated the ALE values associated with the FEF and IFJ clusters in our main results, and potentially also their spatial extent. A second limitation of our study is that although we strived to include as many functional localizers as possible, particularly in the IFJ sample many of the included coordinates were only based on the results of the main fMRI task, and not an independent fMRI localizer. It is not clear to what extent these IFJ activations can be replicated, and there may be a file drawer problem in this sample, which is typically an issue in ALE meta-analyses (Acar et al. 2018). Finally, even though we tried to include experiments that tapped onto similar cognitive functions (i.e., spatial attention and oculomotor control for the FEF, and attention, working memory and cognitive control for the IFJ) to infer spatial convergence within FEF and IFJ, these experiments didn't measure a single cognitive function but rather a collection of several distinct functions and sub-processes, some of which share an important degree of overlap. The detailed investigation of each separate component of these functions/processes, which are often subsumed under the umbrella term of executive functions, would require examining a broader number of paradigms than we included in our analyses, and could lead to further insights into anatomo-functional dissociations in the prefrontal cortex.
